# β-cell responses to high fat feeding: A role and mechanism for redox sensing by SENP1

**DOI:** 10.1101/2022.04.05.487203

**Authors:** Haopeng Lin, Kunimasa Suzuki, Nancy Smith, Xi Li, Lisa Nalbach, Sonia Fuentes, Aliya F Spigelman, Xiaoqing Dai, Austin Bautista, Mourad Ferdaoussi, Saloni Aggarwal, Andrew R Pepper, Leticia P Roma, Emmanuel Ampofo, Wen-hong Li, Patrick E MacDonald

## Abstract

Pancreatic β-cells respond to metabolic stress by upregulating insulin secretion, however the underlying mechanisms remain unclear. In β-cells from overweight humans without diabetes, and mice fed a high-fat diet for 2 days, insulin exocytosis and secretion are enhanced without increased Ca^2+^ influx. β-cell RNA-seq suggests altered metabolic pathways early following HFD, where we find increased basal oxygen consumption, proton leak, but a more reduced cytosolic redox state. Increased β-cell exocytosis after 2-day HFD is dependent on this reduced intracellular redox and requires the <u>sen</u>trin-specific SUMO-<u>p</u>rotease-<u>1</u> (SENP1). Mice with either pancreas- or β-cell-specific SENP1 deletion fail to up-regulate exocytosis and become rapidly glucose intolerant after 2-day HFD. Mechanistically, redox-sensing by SENP1 requires a thiol group at C535 which together with Zn^+^-binding suppresses basal protease activity and unrestrained β-cell exocytosis and increases SENP1 sensitivity to regulation by redox signals.

## Introduction

Type 2 diabetes (T2D) occurs when insulin secretion from pancreatic β-cells fails to meet peripheral demand, which is increased with insulin resistance often coincident with obesity (*1*). While obesity and insulin resistance are major risk factors for T2D, most individuals with obesity maintain normoglycemia as insulin secretion is increased adequately through the up-regulation of β-cell function and mass (*2–4*), although the relative contribution of each remains unclear (*1*). Up-regulation of β-cell insulin secretory capacity precedes increases in islet mass in mice on a high-fat diet (HFD), and at early time points β-cell functional changes likely outweigh structural changes in increased insulin responses (*5, 6*). Loss of β-cell functional up-regulation may contribute to early progression and deterioration to T2D even with intact β-cell mass (*7*).

Upon short-term starvation, insulin secretion is attenuated with a shift from glucose metabolism to fatty acid oxidation (*8*). On the other hand, overnutrition leads to a compensatory increase of insulin secretion with adaptations in β-cell stimulus-secretion coupling. These include enhanced intracellular [Ca^2+^]_i_ responses and up-regulation of metabolic coupling factors that potentiate Ca^2+^ efficacy and amplify insulin secretion (*9–11*). In a longitudinal study measuring parallel β-cell insulin secretion and [Ca^2+^]_i_ responses, the efficacy of Ca^2+^-induced insulin secretion was enhanced via Epac signaling in the early stages of prediabetes (*5*). This may be accompanied by up-regulation of reducing equivalents (*12–14*) and the exocytotic machinery *per se* in prediabetes (*15*), subsequently followed by a reduced expression in T2D (*16*).

SUMOylation is the posttranslational conjugation of small ubiquitin-like modifier (SUMO) peptides to target proteins, which can be reversed by the sentrin-specific proteases (SENPs) such as SENP1. The activity of SENP1 is subject to redox regulation that may involve direct oxidation-reduction of cysteine residues near the enzyme catalytic site (*17*), although the exact structure-function relationship of SENP1 redox-sensing remains unclear. Glucose stimulation of β-cells increases cytosolic reducing signals (*18, 19*) and enhances insulin secretion in part via SENP1, which modifies several exocytotic and exocytosis-related proteins to facilitate insulin granule priming without significantly affecting Ca^2+^ entry (*20, 21*). In a chronic overnutrition model, glucose-stimulated insulin secretion (GSIS) is impaired, in part due to partial inactivation of SENP1 by oxidative stress (*22, 23*). In contrast, it remains unknown whether, at an early compensatory or prediabetic stage, altered redox state and SENP1 activity affectβ-cell functional compensation and glucose homeostasis.

We therefore assessed the role for redox signaling in the up-regulation of β-cell exocytosis that occur early upon high fat feeding in mice, and the structure-function relationship of SENP1 redox sensing. A reduced β-cell redox, commensurate with metabolic rewiring and an increased basal oxygen consumption, enhances exocytotic capacity via a mechanism that requires SENP1. In pancreas- and β-cell–specific SENP1 knockout mice, we demonstrate the requirement for SENP1 in the maintenance of glucose homeostasis within 2 days of high fat feeding via upregulated insulin secretion. Finally, we show that C535 near the catalytic site, together with an interaction with Zn^2+^, is required for redox-regulation of enzyme activity and control of β-cell exocytosis.

## Results

### Higher β-cell secretory capacity from overweight humans and short-term HFD-fed mice

Islets from non-diabetic human donors of 21-45 years of age and BMI >25 exhibited significantly elevated glucose-stimulated insulin secretion (GSIS) compared to similarly aged donors with BMI <25. This greater functionality was absent in T2D (BMI > 25) (**Fig. 1A**). Depolarization-induced β-cell exocytosis is amplified by glucose (*20*). Intriguingly, β-cells from the donors with BMI >25 did not show any difference in depolarization-induced exocytosis compared to BMI <25 at 5- or 10-mM glucose. Instead, the exocytotic response was elevated in the non-diabetic donors with BMI>25 at 1 mM glucose (**Fig. 1B**), suggesting an increased insulin granule priming or amplifying effect even at low glucose. As with insulin secretion, this greater exocytotic capacity at 1 mM glucose was missing in β-cells of donors with T2D and BMI>25 (**Fig. 1B**). These differences were not as clear in islets and β-cells from older donors (age >45), although insulin secretion and β-cell exocytosis were still lower in T2D (**Fig. 1C, D**).

**Figure 1.**
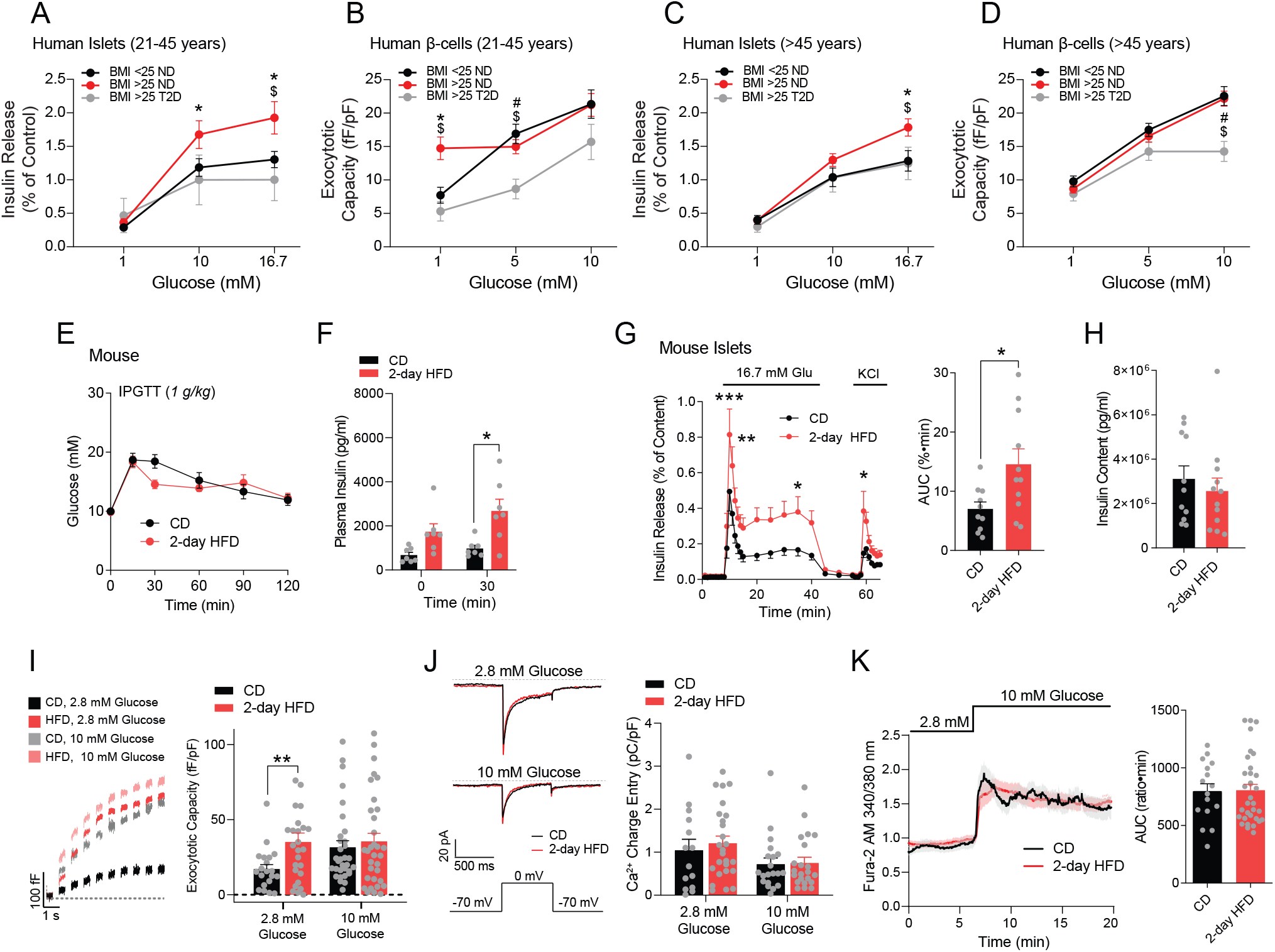
Islets from overweight human donors and 2-day HFD mice exhibited higher insulin secretion and β-cell exocytosis. **(A):** Insulin secretion from islets of young (21-45 years of age) human donors. (N=27-47 non-diabetic donors and 4-5 diabetic donors per condition). **(B):** Exocytotic response of β-cells from young (21-45 years) human donors. (N=11-33 non-diabetic donors with 63-236 cells and 4-5 diabetic donors with 21-36 cells per condition). **(C):** Insulin secretion from islets of older (>45 years of age) human donors. (N=40-111 non-diabetic donors and 18-21 diabetic donors per condition). **(D):** Exocytotic response of β-cells from older (>45 years) human donors. (N=30-73 non-diabetic donors with 164-473 cells and 15-18 diabetic donors with 78-112 cells per condition). **(E):** IPGTT of mice after CD and 2-day HFD (n=14 and 14 mice). **(F):** Plasma insulin level during IPGTT (n=7 and 8 mice). **(G, H):** Insulin secretion (panel G) and content (panel H; n=10 and 11 mice). AUC – area under the curve. **(I):** Representative traces (left), and average total responses, of β-cell exocytosis elicited by a series of 500 ms membrane depolarizations from −70 mV to 0 mV at 2.8- and 10-mM glucose (N=3 pairs of mice, n=21-33 cells). **(J):** Representative traces, and average voltage-dependent Ca^2+^ currents, of β-cell elicited by a single 500 ms membrane depolarization from −70 mV to 0 mV at 2.8- and 10-mM glucose (n=15-26 cells). **(K):** Single cell [Ca^2+^]_i_ response (n=16, 34 cells from 5 and 4 mice). Data are mean ± SEM and were compared with student t-test, one-way or two-way ANOVA followed by Bonferroni post-test. \**P* < 0.05, \*\**P* < 0.01 unless in A-D, \**P* < 0.05 (BMI>25 ND vs BMI<25 ND), $*P* < 0.05 (BMI>25 ND vs BMI>25 T2D), #*P* < 0.01 (BMI<25 ND vs BMI>25 T2D).

To investigate β-cell functional changes that occur early under metabolic stress, we fed male C57BL/6NCrl mice either a chow diet (CD) or HFD for 2 days. Mice fed a 2-day HFD maintained normal glucose tolerance in an intraperitoneal glucose tolerance test (IPGTT) (**Fig. 1E**), along with increased plasma insulin (**Fig. 1F**). GSIS was increased from islets of 2-day HFD mice (**Fig. 1G**), without any effect on insulin content (**Fig. 1H**). Comparatively, after a 4-week HFD, mice were glucose intolerant (**Fig. S1A**) with increased plasma insulin (**Fig. S1B**), increased islet insulin content, and enhanced *in vitro* insulin secretion (**Fig. S1C-E**). In β-cells from 2-day (**Fig. 1I**) and 4-week (**Fig. S1F**) HFD fed mice, exocytotic responses are increased at low glucose, to the same level as at elevated glucose. This suggests that signals which amplify insulin exocytosis, normally upon increased glucose, may be up-regulated even at low glucose. There were no differences in voltage-activated Ca^2+^ entry (**Fig. 1J**) and intracellular Ca^2+^ responses (**Fig. 1K**; **Fig. S1G**), consistent with previous demonstrations of ‘increased Ca^2+^ efficacy’ under short-term HFD (*5*). Indeed, under conditions that ‘clamp’ intracellular Ca^2+^ responses to assess glucose-dependent amplification of insulin secretion (*24*), islets from 4-week HFD fed mice exhibited enhanced KCl-stimulated insulin secretion only at low glucose (**Fig. S1E**).

### Shifts in metabolic pathway expression after short-term HFD

To gain insight into the mechanism contributing to increased insulin secretion following short-term HFD (*25*), we performed RNA sequencing on FACS-sorted β-cells from 2-day CD and HFD mice. We found 213 differentially expressed (DE) genes (**Fig. 2A**). Enrichment analysis identified pathways (PW) such as histone methylation (*26*), MAPK signaling, cholesterol biosynthesis, insulin receptor signaling, and ER-associated misfolded protein response (**Fig. 2B**) (*26, 27*). Histone methylation, the most significantly enriched pathway, can interact or overlap with genes in other enriched pathways, such as MAPK signaling, consistent with the ability of methylation to adapt insulin secretion through MAPK and regulation of glucose metabolism (*26*). All DE genes were submitted to the STRING database (*28*) for protein-protein interaction. Gene set enrichment analysis (GSEA) showed that up-regulated genes were enriched for glycolysis (*Tpi1, Pgk1, Pkm*…) and oxidative phosphorylation (Uqcrq, Vcp, Atp5g3…) pathways (**Fig. 2C**). Interestingly, the cholesterol biosynthesis pathway (*Fdft1, Hsd17b7, Msmo…*) was significantly down-regulated (**Fig. 2C, D**). Cholesterol biosynthesis relies heavily on NADPH consumption (*29*), and its inhibition may enhance insulin secretion by increasing NADPH (*30*). Therefore, the downregulation of cholesterol biosynthesis genes could spare cytosolic NADPH, which might be built up by increased glucose metabolism, for the amplification of insulin release (**Fig. 2D**).

**Figure 2.**
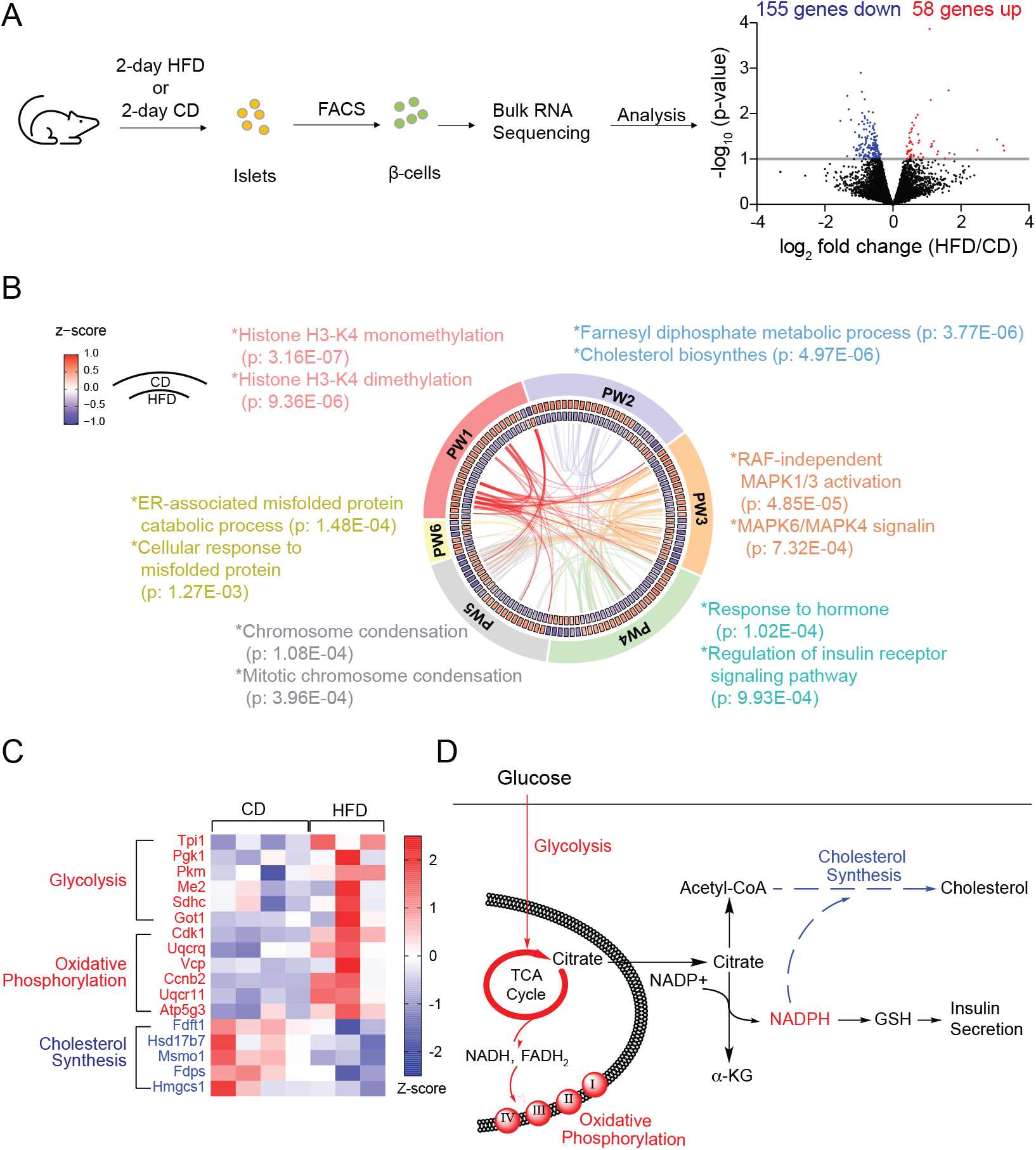
RNA sequencing of purified β-cells following 2-day HFD. **(A):** β-cells from CD and 2-day HFD were isolated through fluorescence activated cell sorting (FACS) for RNA sequencing. 213 genes were identified differentially expressed (DE) genes after 2-day HFD (n=213 genes, N=4, 3 mice). **(B):** All DE genes were submitted to Metascape for functional enrichment analysis. Expression of DE genes on CD and HFD were normalized to z-scores by gene and colorized in circos heat map. Enrichment and cluster analysis were performed on Metascape and the six most enriched pathways (PW) are shown. All DE genes were submitted to STRING database for protein-protein interaction. The related protein interactions for DE genes enriched in the six PW were highlighted as links in the circos plot. **(C):** Gene set enrichment analysis (GSEA) showed significant up- and down-regulated pathways after HFD with false discovery rate (FDR) less than 0.05. All nominal *p*-values were less than 0.01. Normalized enrichment score (NES) reflects the degree to which a gene set is overrepresented in a ranked list of genes. **(D):** Illustration of transcriptomic changes related to metabolism after 2-day HFD.

### Increased β-cell exocytosis after 2-day HFD requires a reducing signal

Glucose metabolism increases insulin exocytosis via a reducing signal (*20*). Functionally, islets from 2-day HFD-fed mice showed higher basal O_2_ consumption and proton-leak (**Fig. 3A,B**). In islets of mice expressing the cytosolic redox sensor Cyto-roGFP2-Orp1(*31*), we found that the cytosol in β-cells from 2-day HFD-fed mice was more reduced compared to control mice fed CD (**Fig. 3C**). A reduced redox state appears required for enhanced β-cell exocytosis since this was recapitulated in β-cells of CD-fed mice upon direct intracellular dialysis of reduced glutathione (GSH), which could not increase exocytosis further in β-cells of HFD-fed mice (**Fig. 3D**) and was reversed by direct intracellular application of H_2_O_2_ (**Fig. 3E**). Previously we showed that SENP1 couples redox state to insulin exocytosis (**Fig. 3F**) (*20*). Our transcriptomic analysis indicated an upregulation of *Senp1*, but not other isoforms, in β-cells after 2-day HFD (not shown), which we confirmed by qPCR (**Fig. 3G**). Direct intracellular dialysis of active SENP1 catalytic domain increases exocytosis in β-cells from CD-fed mice but, like GSH, cannot increase exocytosis further in the β-cells of HFD-fed mice (**Fig. 3H**).

**Figure 3.**
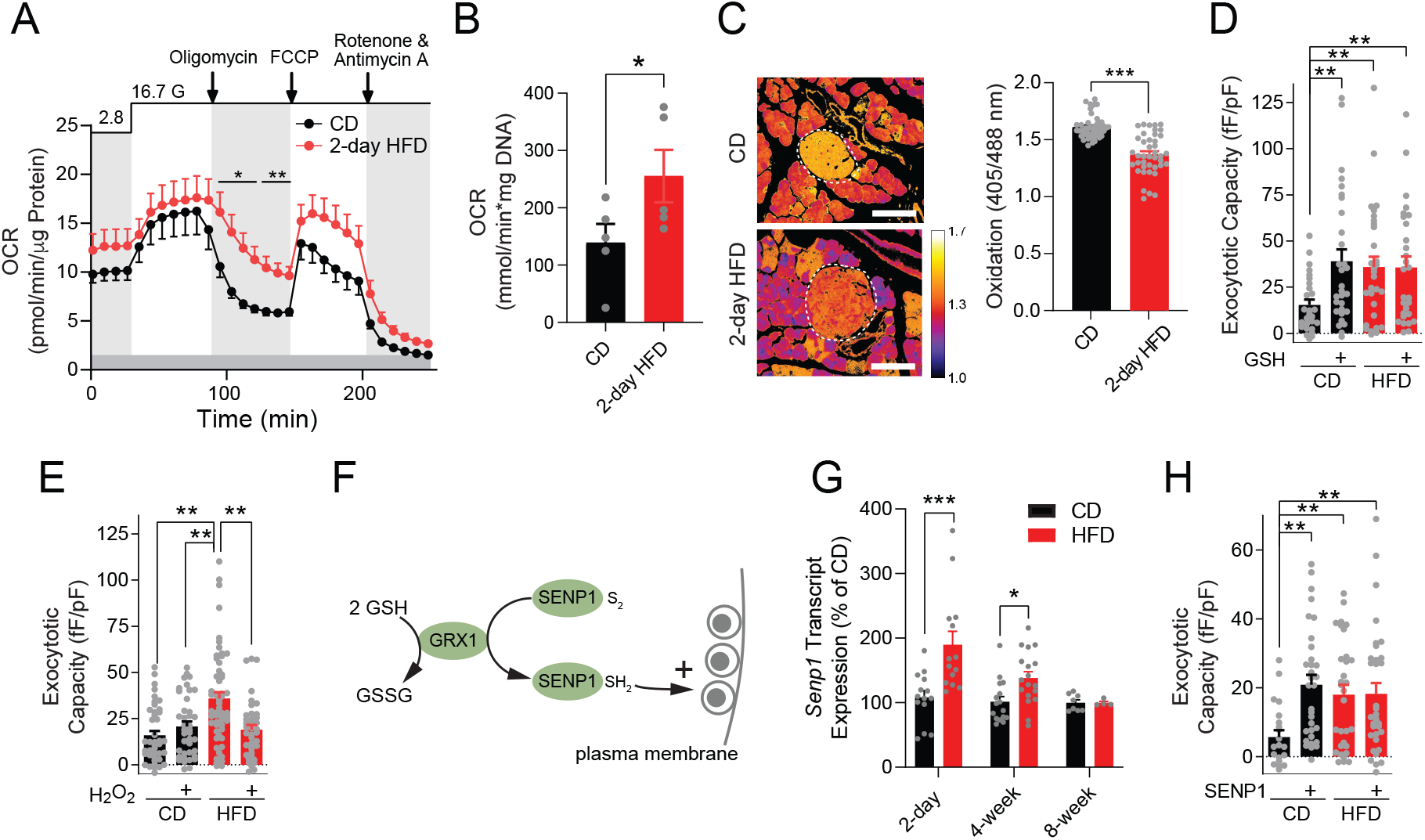
Reductive cytosolic redox signaling via SENP1 contributes to enhanced exocytosis during 2-day HFD. **(A):** Oxygen consumption rate (OCR) measured by Seahorse assay (N=6, 6 mice). **(B)**: OCR measured by Fluorescence Lifetime Micro Oxygen Monitoring at 2.8 mM glucose (N=4, 5 mice). **(C):** Representative image of pancreatic islets (white dashed circle) carrying cyto-roGFP2-Orp1 sensor (left). Scale bar = 100 µm. Redox ratio of individual islets (n= 65, 41 islets from 6 pairs of mice). **(D, E):** Exocytosis with infusion of (panel D) 10 µM GSH or (panel E) 200 µM H_2_O_2_ at 2.8 mM glucose after 2-day HFD. (C - n=26-56 cells from 4, 4 mice, D - n= 39-56 cells from 7, 7 mice). **(F):** Illustration of redox-control of insulin exocytosis. **(G):** *Senp1* expression by qRT-PCR in islets from CD or HFD fed mice (N=2-9 mice). **(H):** Exocytosis with infusion of 4 μg/mL catalytic SENP1 (n=21-33cells from 4, 4 mice per group) at 2.8 mM glucose. Data are mean ± SEM and were compared with student t-test, one-way ANOVA followed by Bonferroni post-test. \**P* < 0.05, \*\**P* < 0.01, * \*\**P* < 0.001.

In pSENP1-KO mice (*22*), males become more rapidly glucose intolerant than littermate controls, with significantly IP and oral glucose intolerance after 2-day HFD (**Fig. 4A-D**). Loss of β-cell SENP1 prevents the up-regulation of exocytosis after either a 2-day (**Fig. 4E**) or 4-week (**Fig. 4F**) HFD. Similar to the pSENP1-KO line, male βSENP1-KO mice (*22*) become rapidly intolerant of IP glucose compared with control littermates upon 2-day HFD (**Fig. 4G, H**), with an impaired plasma insulin response (**Fig. 4I**). These responses were less obvious in female mice from both the pSENP1-KO and βSENP1-KO lines (**Fig. S2, S3**), which are more resistant to the development of insulin resistance upon HFD (*32*). Also, after 4-week HFD, oral glucose intolerance became more prominent in the βSENP1-KO females **(Fig. S3I**) and males (**Fig. 4J-L, Fig. S4**), although again the females were generally more insensitive, consistent with our recent report at 8 weeks of HFD in both the pSENP1- and βSENP1-KO models (*22*).

**Figure 4.**
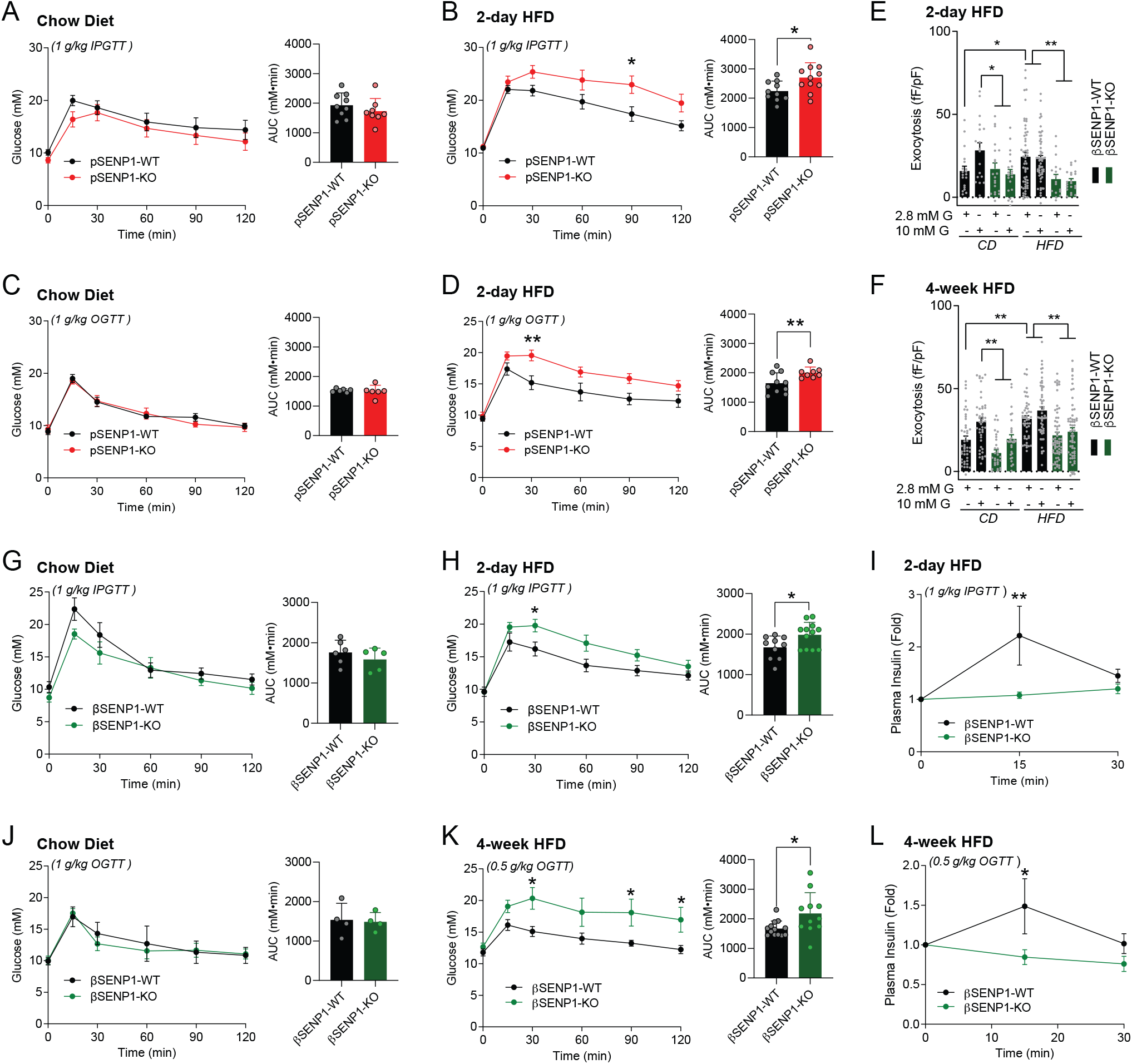
βSENP1-KO mice develop glucose intolerance following 2-day HFD. **(A-B):** IPGTT of male pSENP1-KO and pSENP1-WT mice fed with CD and 2-HFD (A - n=9, 8 mice, B - n=10, 11). **(C-D):** OGTT of male pSENP1-KO and pSENP1-WT mice fed with CD and 2-HFD (C - n=6, 6 mice, D - n=10, 8). **(E-F):** Exocytosis from β-cells of βSENP1-WT and βSENP1-KO mice after 2-day and 4-week HFD (A - n=18-56 cells, B - n=30-65 cells). **(G-H):** IPGTT of male βSENP1- KO and βSENP1-WT mice fed with CD and 2-day HFD (G - n=6, 5, H - n= 10, 12). **(I):** Plasma insulin levels during IPGTT after CD or 2-day HFD (n=8, 8). **(J-K):** OGTT of male βSENP1-KO and βSENP1-WT mice fed with CD and 4-week HFD (J - n=4, 4, K - n= 13, 11). **(L):** Plasma insulin levels during OGTT after CD or 4-week HFD (n= 8, 8). Data are mean ± SEM and were compared with student t-test or two-way ANOVA followed by Bonferroni post-test. \**P* < 0.05, \*\**P* < 0.01.

### Redox regulation of exocytosis requires SENP1 C535

SENP1, a cysteine protease, is subject to redox regulation via a thiol group on the catalytic cysteine C603 or possibly other cysteines (*17*). C603 is the most conserved cysteine across SENP isoforms and species, while C535 is the least conserved near the active site (**Fig. 5A**). Although redox regulation of intra-molecular disulfide bonds may control cysteine protease activity, as in GRX1 and TRX1 (*33*), the key cysteines of the SENP1 catalytic domain appear too distant from the catalytic C603 to facilitate such regulation (*34*) (**Fig. S5A**). We therefore generated a series of recombinant SENP1 catalytic domain proteins with cysteine-to-serine substitutions and measured SUMO-protease activity *in vitro* (**Fig. S5B**). While mutation of the catalytic cystine (C603S) completely abolished SENP1 activity as expected, the C535S mutations maintained a 2-fold higher activity compared to the SENP1 WT and other mutants (**Fig. 5B**). Under conditions where SENP1 is fully activated (minimal oxidization and absence of divalent cations), the activities of SENP1 WT and SENP1 C535S were similar, and while the WT was strongly inhibited by 5 mM H_2_O_2_, the C535S was not (**Fig. 5C**). Intracellular dialysis of either SENP1 WT or C535S into human β-cells increased exocytotic capacity (similar to that in young, overweight, donors). While co-dialysis of H_2_O_2_ blocked this action of SENP1 WT, the C535S mutant maintained its functional activity (**Fig. 5D**). The activity of SENP1 WT could be increased by GSH together with GRX1, while SENP1 C535S appeared maximally activated (**Fig. 5E**). Altogether these results suggest that C535 is required for SENP1 sensitivity to redox and its function in augmenting insulin exocytosis.

**Figure 5.**
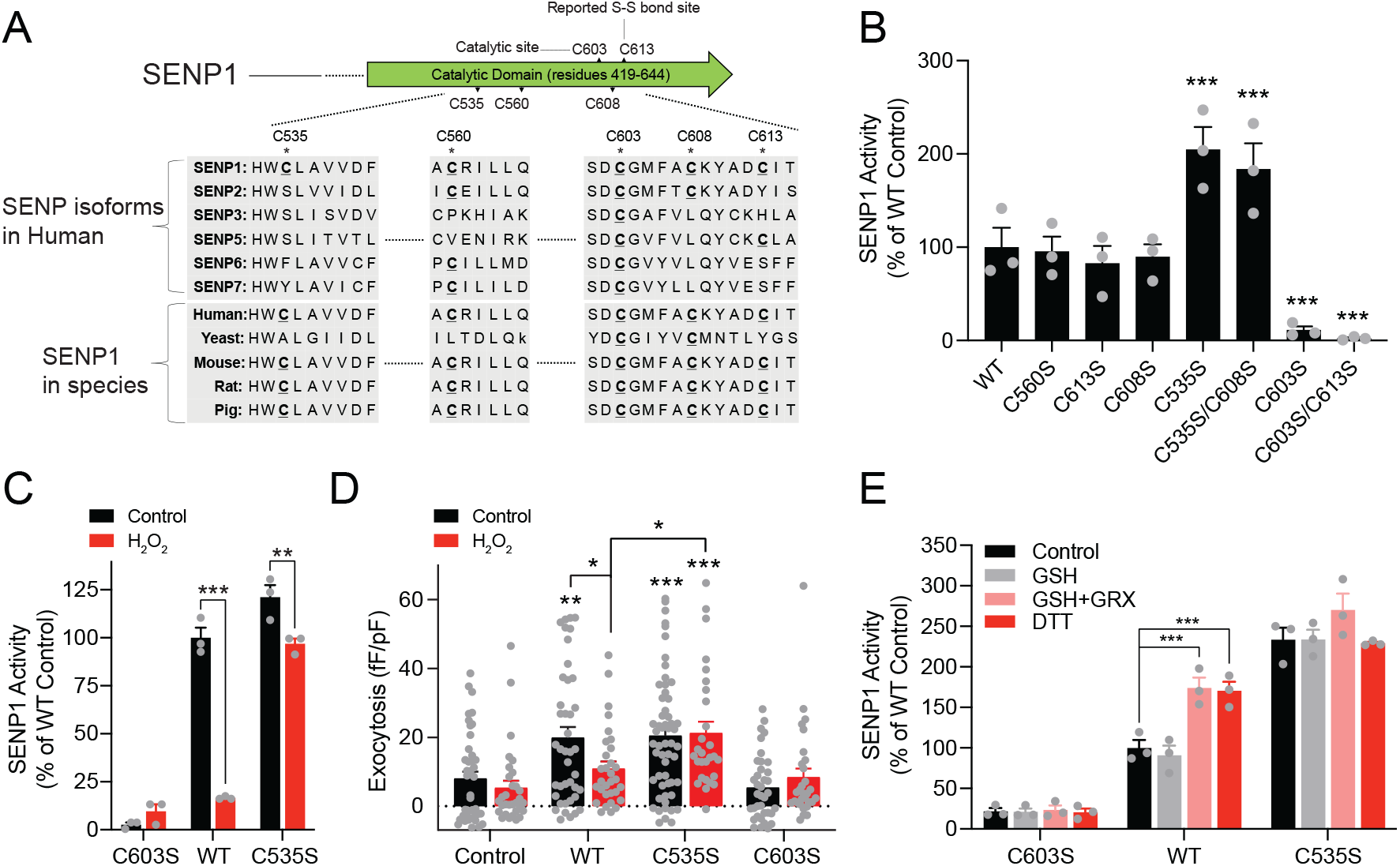
Redox regulation of SENP1 activity requires C535. **(A):** Multiple amino acid comparison among different isoforms of SENP across species. **(B):** SENP1 activity after indicated cysteine-to-serine substitution (n=3). In hindsight, some recombinant enzymes were likely partially inactivated by oxidization upon prolonged exposure to air. **(C):** Activity of SENP1 C603S, WT, and C535S upon full activation by DTT (10 mM), and subsequent inhibition by 5 mM H_2_O_2_ (n=3). **(D):** 4 μg/mL glutathione-S-transferase (GST) peptide (Control), SENP1 WT, C535S and C603S were infused into human β-cells with/without 200 µM H_2_O_2_ to assess their effects on exocytosis at 2.8 mM glucose (n=28-46 cells per group from 7 human donors). **(E):** Activity of SENP1 C603S, WT, and C535S (following partial inactivation by oxidization upon exposure air) in the presence of GSH (0.1 mM) along or with GRX1 (10 µg/ml). DTT (10 mM) was used to fully activate the enzymes (n=3). Data are mean ± SEM and were compared with student t-test, one-way ANOVA or two-way ANOVA followed by Bonferroni post-test. \**P* < 0.05, \*\**P* < 0.01, \*\*\**P* < 0.001. In panel D, * indicated comparison between SENP1 WT and C535S.

We wondered whether an additional mechanism contributes to suppression of baseline SENP1 activity. Zn^2+^-cysteine is a critical mediator of redox-regulation and protease activity (*35*). *In silico* analysis predicts that C603-C535-H533 is a Zn^2+^-binding site in SENP1 (**Fig. S5C**), and we found that Zn^2+^, and to a lesser extent Ni^2+^ (present during re-folding in some of our earlier experiments) inhibited the activity of SENP1 WT but not SENP1 C535S (**Fig. 6A**). The IC_50_ for SENP1 inhibition by Zn^2+^ is right-shifted by nearly-30-fold in the C535S mutant, indicative of allosteric interaction of C535 with Zn^2+^ to inhibit SENP1(**Fig. 6B**). This can be readily reversed by chelation with EDTA (**Fig. 6C**) and was not due to non-specific effect of Zn^2+^ on His×6 tag from recombinant SENP1 (**Fig. S5D**).

**Figure 6.**
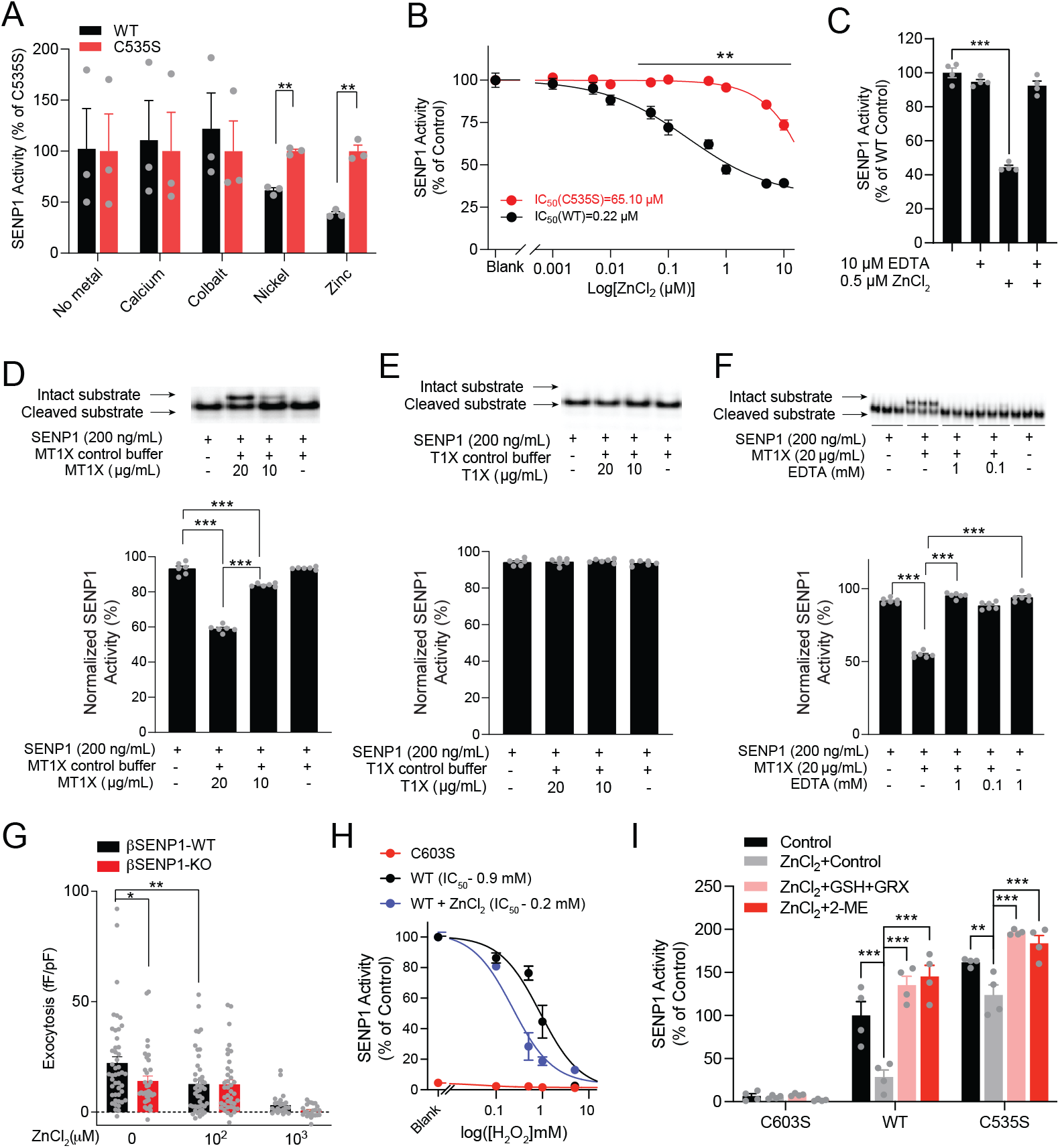
Zn^2+^ tunes SENP1 redox sensitivity and SENP1-dependent β-cell exocytosis. **(A):** Activity of SENP1 WT and C535S in the presence or absence of Ca^2+^, Co^2+^, Ni^2+^ and Zn^2+^ (n=3). Data were normalized to C535S activity, as this is more resistant to oxidation (n=3). **(B):** Dose-response curve of SENP1 inhibition by ZnCl_2_ (n=3). **(C):** SENP1 inhibition by ZnCl_2_ could be reversed with the chelating reagent EDTA. **(D-F):** SENP1 activity, measured using a native-PAGE assay, inhibited by Zn^2+^-carrying metallothionine (MT1X; panel D) but not Zn^2+^-depleted thionine (T1X; panel E) or by MT1X in the presence of EDTA (panel F) (n=6). **(G):** Effect of Zn^2+^ on exocytosis from b-cells of βSENP1-WT and -KO mice at 5 mM glucose (n= 23-46 cells from 6 pairs of mice). **(H):** Concentration-response curve of SENP1 activity, with enzyme prepared with or without ZnCl_2_ during refolding, to inhibition by H_2_O_2_. Activity of SENP1 C603S is shown for comparison (n=3-6). **(I):** Activity of SENP1 C603S, WT, and C535S in the presence of 1 µM Zn^2+^ and subsequent activation with 5 mM GSH and 10 µg/ml GRX1. 2-mercaptoethanol (2-ME, 1 mM) was used to fully activate enzymes (n=4). Data are mean ± SEM and were compared with one-way ANOVA or two-way ANOVA followed by Bonferroni post-test. \**P* < 0.05, \*\**P* < 0.01, \*\*\**P* < 0.001.

While Zn^2+^ inhibits SENP1 WT with an IC_50_ (0.22 uM), similar to the well-established Zn^2+^-inhibition of enzymes (*36*) including proteases such as the caspases (IC_50_ = 0.1-3.4 µM; (*37, 38*)), this is above the free cytoplasmic Zn^2+^ (sub-nM) in eukaryotic cells (*39*). Compared to the *in vitro* Zn^2+^ titration assay, regulation of Zn^2+^ activity in a cellular environment is more complex, yielding compartmentalized Zn^2+^ signaling events with defined spatiotemporal characteristics. We examined the potential involvement of metallothionein, a Zn^2+^-binding protein participating in Zn-storage and distribution to other proteins (*40*). Metallothionein is abundantly present in the pancreas (average 264 µg/mL or 43.6 µM in human pancreas (*41*). Because MT1X interfered with the binding of Hisx6 tagged SENP1 substrate (Hisx6-SUMO-mCherry) to Nickel-NTA agarose for the SENP1 assay, (see Methods), we developed an alternative assay that does not require Nickel-NTA agarose (**Fig. S5D**). Metallothionein 1X (MT1X), a subtype present in β-cells, negatively regulates insulin secretion (*42*) and suppresses SENP1 activity similarly to free Zn^2+^ (**Fig. 6D**), while thionein 1 (T1X; an apoprotein of MT1X not carrying Zn^2+^) does not (**Fig. 6E**). Zn^2+^ chelation with EDTA also reverses SENP1 inhibition by MT1X (**Fig 6F**). In β-cells of the βSENP-WT mice, dialysis of 100 µM ZnCl_2_ impaired exocytosis without decreasing Ca^2+^ currents, to a level similar to that in the βSENP-KO (**Fig. 6G; Fig. S6A**). While 100 µM ZnCl_2_ did not decrease exocytosis further in the βSENP-KO, higher concentrations of ZnCl_2_ (1 mM) dramatically decreased exocytosis in both groups possibly by inhibiting Ca^2+^ currents directly (*43*) (**Fig 6G, Fig. S6A**). Finally, in the presence of Zn^2+^ SENP1 was more sensitive to inactivation by H_2_O_2_ (**Fig. 6H**), and more robustly activated by GSH and GRX1 (5.3-fold) while the C535S mutant remained more active (**Fig. 6I**).

## Discussion

β-cell compensation and decompensation is a determinant of early T2D progression (*44*), yet the underlying mechanisms remain unknown. Here we demonstrate that a reducing signal requiring SENP1 facilitates an upregulation of β-cell function to maintain glucose tolerance very early after high fat feeding. Redox-sensing by SENP1 relies on C535 and is tuned by an interaction with Zn^2+^. Notably, the enhanced depolarization-induced exocytosis observed at low glucose soon after 2-day HFD is recapitulated by intracellular dialysis of reducing signals or active SENP1, which exerts a similar priming effect (*21*), and is lost upon β-cell knockout of SENP1. This is not expected to increase basal insulin secretion in the absence of an increased intracellular Ca^2+^ concentration. Instead, an increased insulin granule priming at low glucose will increase the size of the releasable pool of insulin granules on which subsequent glucose-dependent Ca^2+^ responses can act, resulting in increased glucose-stimulated insulin secretion. This is consistent with the enhanced efficacy of Ca^2+^ in 1-week HFD-fed mice (*5*) and pre-diabetic *db/db* mice (*15*).

Redox signals are important regulators of insulin secretion and functional compensation (*12, 20*), although there is debate as to whether reactive oxygen species (*45*) or reducing signals (*46*) augment insulin secretion. Possibly both are true, depending on timing, localization, and mechanisms involved, such as the modulation of excitability by reactive oxygen species *via* K^+^ channels (*47*) or by the direct regulation of membrane fusion by reducing signals (*48*). We observed a remodeling of metabolic pathways in RNAseq data and increased proton leak and basal O_2_ consumption after 2-day HFD. Coincident with this, we observe a more reduced cytosol in the β-cell. Although the underlying mechanism remains unclear, down-regulation of cholesterol biosynthesis as seen in the RNAseq data may contribute to an increase in cytosolic NADPH (*30*). Increased mitochondrial proton-leak may also contribute to higher insulin secretion (*49*), and increased sub-maximal glucose-fueled mitochondrial TCA cycle flux may generate higher cytosolic reducing equivalents (*50*). We and others have shown that NADPH, produced by either the mitochondrial export of reducing equivalents or the pentose phosphate pathway (*51, 52*), augments Ca^2+^-triggered insulin exocytosis (*20, 48, 53*). One caveat is that the Cyto-roGFP2-Orp1 redox-sensor reports the steady state balance between oxidizing and reducing signals (*19, 54*). The reduced redox could result from either decreased H_2_O_2_ production or increased reducing equivalents. Adaptive glucose metabolism may also enhance *de novo* GSH biosynthesis independent of NADPH generation (*55*). Therefore, increased *de novo* GSH biosynthesis could contribute to a compensatory increase in insulin secretion. Nonetheless, a reduced cytosolic redox state appears critical for the increased β-cell exocytosis seen after 2-day HFD since it could be reversed by H_2_O_2_ and replicated by GSH or NADPH.

Compared to other isoforms, SENP1 is a redox-sensitive protease with an unique interaction between a free thiol on the catalytic cysteine (C603) and other cysteines (*17*). C613 is reported to be a ‘positive’ redox sensor by forming an intermolecular disulfide bond with the C603 and protect SENP1 from irreversible oxidation (*17*). We find however that mutation of C613 had little effect on SENP1 activity, but rather that C535 appears to restrain SUMO-protease activity. The C535S mutant had 2-fold higher activity and was resistant to inactivation by H_2_O_2_, indicating that C535 is a ‘negative’ redox sensor. Indeed, the atomic distance between thiols of C535 and C603 prevents formation of an intramolecular disulfide bond, consistent with the absence of disulfide bonds in available SENP1 crystal structures (*34*). We also find that C535 does not control enzyme activity via the formation of intermolecular SENP1 dimers (not shown). Thus, it seems most likely that SENP1 redox-sensing via C535 occurs via a proton transfer pathway (**Fig. S6B**), and deprotonation of the thiol group on C603 to activate SENP1. Specifically, proton transfers from the thiol group of C603 to the carboxyl group of D550 via H533 results in the conversion of a thiol on C603 to an active thiolate in SENP1 (*56*). C535 may donate a proton to either H533 or D550, like C603 does, to competitively restrain proton transfer from C603, leading to inhibition of SENP1, under oxidizing conditions.

SENP1 couples reducing power to insulin secretion (*20, 21, 57, 58*). Interestingly, we find that Zn^2+^ allosterically inhibits SENP1 in a manner that is dependent upon C535. Mechanistically, the predicted Zn^2+^-binding site includes C603, H533 and C535, which is one of the most ubiquitous Zn^2+^ binding motifs (*59*). A coordinate bond between Zn^2+^ and these residues likely further restricts proton transfer. Zn^2+^ coordination is expected to lower the pKa of a thiol by two orders of magnitude, facilitating the formation of a thiolate (*60*). Indeed, thiol-binding Zn^2+^ readily dissociates in the presence of oxidant, leaving a highly reactive thiolate for oxidant (*61*). Therefore, interaction between C535 and Zn^2+^ sensitizes SENP1 to oxidative inhibition and allows robust reactivation by reducing signals, not dissimilar to the reported interaction between reactive oxygen species and Zn^2+^ in the regulation of protein tyrosine phosphatases (*62*).

Zn^2+^ inhibits exocytosis via SENP1 with a calculated free Zn^2+^ at ∼0.08 µM. However, cytosolic free Zn^2+^ levels are not perturbed by glucose stimulation (*63*) or the short-term HFD (not shown). Zn^2+^ binding of SENP1 may occur at the time of protein folding in the ER, or via direct interaction with Zn^2+^-binding/buffering proteins such as MT1X which have been shown to control islet Zn^2+^ fluctuations (*64*) and inhibit glucose-stimulated insulin secretion via a so-far unidentified interaction with the exocytotic machinery (*42*). While free Zn^2+^ is in the low-nM-range in the β-cell cytosol (*63*), the abundant Zn^2+^-binding MT1X likely transfers Zn^2+^ to SENP1, tuning its redox-sensitivity and activity. During β-cell compensation, MT1X was downregulated to increase insulin secretion (*42*). Consistent with this our transcriptomic data showed that expression of *Mt1* was decreased after 2-day HFD (not shown). The combined effect of less Zn^2+^ transferred to SENP1, upregulation of *Senp1* expression, and a reduced redox state would be expected result in a more active SENP1 and augmentation of β-cell exocytosis.

Knockout of islet and β-cell SENP1, in two separate Cre-driver models, led to IP glucose intolerance after 2-day HFD associated with a loss of insulin response. However, oral glucose tolerance was not different following 2-day HFD in most cases and in contrast with the 4-week HFD where the SENP1-KO showed selectively impaired oral, not IP, glucose tolerance. This discrepancy highlights the adaptive changes that occur during different stages of high fat feeding (*65*). The incretin response is paramount in limiting oral glucose intolerance after long-term HFD (*22, 66*), where incretin release from the gut (*22*) and β-cell incretin receptors (*65*) are upregulated. β-cell SENP1 is required for these compensational changes to maintain incretin-stimulated insulin secretion and oral glucose tolerance (*22, 67*). However, the β-cell GLP-1 receptor is not necessary for oral glucose tolerance under normal conditions (*68*), similar to the role of SENP1 for oral glucose tolerance under the same conditions (*22*) or after 2-day HFD, under which circulating incretins remained low (not shown). Additionally, the extra-islet action of GLP-1 could mask the effect of β-cell SENP1, such that OGTT is similar in βSENP1-KO and -WT mice (*68*). Finally, we note that the less cell-type-selective pSENP1-KO mice, which have impaired GIP-stimulated glucagon secretion (*22*), did show impaired oral glucose tolerance after 2-day HFD, indicating the potential importance of α-to-β cell communication under this condition and as previously reported (*66*).

Thus, we show that insulin secretion is upregulated from islets from human donors with increased BMI, at early/mid-age, and from mouse islets shortly after high fat feeding. This is not due to an increased Ca^2+^ response in β-cells, but to an increase in the availability of releasable insulin granules that is promoted at low glucose (i.e., prior to glucose-stimulation). This requires a reducing signal, mediated by the SUMO-protease SENP1 via a sensing mechanism involving a Zn^2+^-dependent redox control of enzyme activity by C535 near the enzyme catalytic site. In mice this enhanced insulin secretory capacity acts to maintain glucose tolerance in the early stages of high fat feeding.

## Materials and Methods

### Islet isolation and cell culture

Human islets were isolated at the Alberta Diabetes Institute IsletCore (www.isletcore.ca) and cultured overnight in DMEM (11885, Gibco) supplemented with L-glutamine, 110 mg/l sodium pyruvate, 10 % FBS (12483, Gibco), and 100 U/ml penicillin/streptomycin (15140, Gibco) (*69*). A total of 265 non-diabetic (ND) donors and 34 T2D donors (HbA1c>6.5) (**Table S1**) were examined. Separation of donors by BMI and age was based on clinical diagnostic criteria for overweight, and previous studies by us (*69*) and others (*70*) demonstrating declines in human islet insulin secretion between the ages of 40 and 50. All human islet studies were approved by the Human Research Ethics Board (Pro00013094; Pro00001754) at the University of Alberta and all families of organ donors provided written informed consent.

C57BL/6NCrl mice (Charles River Laboratories) were fed standard chow diet (5L0D*, PicoLab Laboratory Rodent Diet) or 60 % high-fat diet (S3282, Bio-Serv) at 12 weeks of age for 2-days or 4-weeks. Pdx1-Cre mice (B6.FVB-Tg (Pdx1-Cre)6 Tuv/J, Jackson Lab, 014647) on a C57BL/6N background and Ins1-Cre mice on a mixed C57BL/6J and SV129 background (*71*) were crossed with *Senp1*-floxed mice on a C57BL/6J background to generate gut/pancreas specific (Pdx1-Cre^+^;SENP1^fl/fl^ - pSENP1-KO) and β-cell-specific knockouts (Ins1-Cre^+^;SENP1^fl/fl^ - βSENP1-KO) (*20, 72*). Control littermates Pdx1-Cre^+^;SENP1^+/+^ mice (pSENP1-WT) and Ins1-Cre^+^;SENP1^+/+^ mice (βSENP1-WT) at 12 weeks of age were used for experiments. Genotypes and SENP1 expression were confirmed as previously (*22*). Mouse islets were isolated by collagenase digestion and purified by histopaque density gradient centrifugation before hand-picking (*22*). All mouse islets or single cells were cultured in RPMI-1640 (11875, Gibco) with 11.1 mM glucose, 10 % FBS (12483, Gibco), 100 units/mL penicillin, and 100 µg/mL streptomycin (15140, Gibco). All studies with mice were approved by the Animal Policy and Welfare Committee (AUP00000291) at the University of Alberta.

### Insulin secretion

After overnight culture, static insulin secretion was measured as previously (*69*) at the glucose concentrations indicated. Diazoxide and KCl concentration were 100 μM and 30 mM respectively. For dynamic insulin responses 25 islets were pre-perifused for 30 minutes at 2.8 mM glucose followed by 16.7 mM glucose and KCl (30 mM) at a sample collection rate of 100 µl/min every 2-5 minutes (*22*). Insulin content was extracted with acid/ethanol. The samples were stored at −20°C and assayed by using Insulin Detection Kit (STELLUX® Chemi Rodent Insulin ELISA kit; STELLUX® Chemi Human Insulin ELISA kit, Alpco).

### Exocytosis measurements

Human and mouse islets were dispersed to single cells in Ca^2+^-free buffer and cultured in 5.5 mM glucose DMEM and 11 mM glucose RPMI, respectively as above. After overnight culture, dispersed cells were preincubated at 1 mM (human) or 2.8 mM (mouse) glucose for 1 hour and patched in bath solution containing (in mM): 118 NaCl, 5.6 KCl, 20 TEA, 1.2 MgCl_2_, 2.6 CaCl_2_, 5 HEPES at different glucose conditions (1, 2.8, 5, 10 mM) with a pH of 7.4 (adjusted by NaOH) at 32-35°C. Whole-cell capacitance was recorded with the sine+DC lock-in function of an EPC10 amplifier and PatchMaster software (HEKA Electronics). Exocytotic responses and inward Ca^2+^ currents were measured 1-2 minutes after obtaining the whole-cell configuration in response to ten 500 ms depolarizations to 0 mV from a holding potential of −70 mV. Changes in capacitance and integrated Ca^2+^ charge entry were normalized to cell size (fF/pF and pC/pF, respectively). The intracellular solution contained (in mM): 125 Cs-Glutamate, 10 CsCl, 10 NaCl, 1 MgCl_2_, 5 HEPES, 0.05 EGTA, 3 MgATP and 0.1 cAMP with pH=7.15 (pH adjusted with CsOH).

In the intracellular dialysis experiments, measurements were 2-3 minutes after obtaining the whole-cell configuration. Compounds or recombinant enzymes were added to pipette solution as indicated. For mouse β cells, these were 200 μM H_2_O_2_ (Sigma), 10 μM reduced glutathione (GSH, Sigma), 4 μg/ml cSENP1 (Enzo Life Technologies) or glutathione-S-transferase (GST) (Enzo Life Technologies) as a control protein. For human β-cells, these were 4 μg/ml GST, wild type SENP1, C535S and C603 mutant recombinant proteins (see below) pretreated with or without 200 μM H_2_O_2_ for 10 minutes. Human β-cells were identified by immunostaining for insulin. Mouse β-cells were identified by size (>4 pF) and a Na^+^ channel half-maximal inactivation at around −90 mV (*73*).

### Glucose homeostasis

Oral glucose tolerance test (OGTT), intraperitoneal tolerance test (IPGTT) and insulin tolerance test (ITT) were performed as previously described (*22*). For OGTT and IPGTT on chow diet or 2-day HFD, glucose concentration was 1 g/kg dextrose, while on 4-week HFD, it was 0.5 g/kg dextrose. Tail blood was collected at indicated times for insulin (STELLUX® Chemi Rodent Insulin ELISA kit, Alpco) and glucose measurement. ITT was performed with a concentration of 1 U/kg Humulin R (Eli Lilly).

### Intracellular Ca^2+^ imaging

Mouse islet cells were cultured on 35 mm glass bottom dishes with 10 mm micro-well (Cellvis) overnight in 11 mM glucose RPMI media (11875, Thermo Fisher). Cells were pre-incubated with Fura-2AM (1 μM) for 15 min and perifused with KRB solution at indicated glucose level and imaged at 0.5 Hz. Fluorescence signal was excited at 340/380 nm (intensity ratio 20:8) and detected at an emission light of 510 nm using Life Acquisition software (Till Photonics) on an inverted microscope (Zeiss Axioobserver, Carl Zeiss Canada Ltd.) equipped with a rapid-switching light source (Oligochrome; Till Photonics, Grafelfing, Germany). β-cells were marked and identified by immunostaining and fluorescence ratios were calculated using ImageJ (NIH).

### Fluorescence activated cell sorting (FACS) of β-cells and mRNA sequencing

Islet α- and β-cells were identified by ZIGIR vs Ex4-Cy5 through FACS analysis as above (*74*). Total RNA was isolated from sorted β-cells using TRIzol reagent (Invitrogen) according to the manufacturer’s protocol and cDNA libraries were prepared using Illumina’s TruSeq Stranded mRNA Library Prep kit. The cDNA libraries were validated by TapeStation DNA 1000 High Sensitivity assay (Agilent) and quantified by Qubit dsDNA High Sensitivity assay (Invitrogen). High quality libraries were submitted to the UT Southwestern Next Generation Sequencing Core facility and RNA sequencing was performed using Illumina’s NextSeq 500 High Output instrument. Raw reads were processed to transcripts per million (TPM) and differentially expressed (DE) genes were initially determined by using the edgeR package. Genes with P-value<0.1, coefficient of variation (CV) of less than 1, same change of direction (either positive or negative) and specific gene counts range (mean gene counts in at least one experimental group should be higher than 5 RPKM) were considered differentially expressed between CD and HFD. All differentially expressed genes are used for downstream analyses and listed in **Data S1**.

Differentially expressed gene function and enriched pathways were identified with Metascape (http://metascape.org/). KEGG Pathway, GO Biological Processes, Reactome Gene Sets, CORUM, TRRUST and PaGenBase were included in pathway enrichment analysis with a p-value <0.01. Terms with an enrichment factor > 1.5 and a similarity of >0.3 are grouped into a cluster represented by the most significant term (*75*). To examine protein-protein interaction (PPI), STRING database (https://string-db.org/) was used and only PPIs with interaction score higher than 0.70 were retrieved and linked in the circus plot by using R-studio (circlize package) (*28, 76*). To run an unbiased gene set enrichment analysis (GSEA) (*77*), all Genes with CV of less than 1, same change of direction and specific gene counts range (at least one experimental group should be higher than 5 RPKM) were submitted and listed in **Data S2**. H.all.v7.4.symbols.gmt[Hallmarks] gene dataset was used as reference data set. Enriched gene sets were considered significant with gene size over 50, false discovery rate less than 0.05, nominal *p*-value less than 0.01.

### Redox measurement

Measurements of the intracellular H_2_O_2_ levels were made using a redox histology approach, as previously described (*31, 78, 79*). Briefly, batches of 30 islets were collected and immersed in 50 mM N-ethyl-maleimide (NEM) dissolved in PBS for 20 min for sensor chemical fixation. The NEM was then removed, and islets fixed in 4 % paraformaldehyde for 30 min. After paraformaldehyde removal, the islets were incubated in 100-µL HepatoQuick (Roth, Karlsruhe, Germany) mixed with human citrate plasma (1:2 v/v) and 1 % CaCl_2_ for 1 h at 37^◦^C. Clots were then incubated in 95 % ethanol at 4 °C overnight and subsequently dehydrated in ethanol prior to paraffin embedding. The paraffin embedded clots were cut in 3 µm thick sections with a semi-automated rotary microtome (Leica Biosystems, Wetzlar, Germany) and placed on silanized glass slides. Images of all the islets in each slide were obtained by the Axio Observer 7 fluorescence microscope (Zeiss, Oberkochen, Germany) with a 20× objective using excitation 405 nm and 488 nm, emission 500-530 nm. The images were analyzed with ImageJ Fiji Software, and the ratio (405/488 nm) was used to compare different groups.

### Oxygen consumption

Agilent’s seahorse XFE24 Analyzer with the islet capture microplates was used and detailed procedure was performed as described (*80*). Briefly, 70 islets per well were assayed. Islets were sequentially treated with 2.8 mM glucose, 16.7 mM glucose, 5 µM oligomycin, 5 µM FCCP and 5 µM rotenone/antimycin A. Each experiment was run with a 3-minute mix, 2-minute wait and 3-minute measurement period for all data points. Each biological replicate had three technical replicates and oxygen consumption rate was normalized to islet protein.

For oxygen consumption measured with the Fluorescence Lifetime Micro Oxygen Monitoring System (FOL/C3T175P, Instech Laboratories Inc.) around 200 islets incubated overnight were washed 3x with serum-free RPMI media at 2.8 mM glucose, added to the chamber for data collection for 20 minutes, and then collected for normalization of oxygen consumption to DNA content measured using a Quant-iT™ PicoGreen™ dsDNA Assay Kits (Invitrogen).

### Preparation of recombinant wild-type and mutant SENP1

Human SENP1 catalytic domain cDNA (NM_001267594/NP_001254523.1, 419-644 residues) was cloned in the downstream of a His×6 coding sequence, between Xba-I and BamH-I sites of the pT7JLH plasmid to add a His×6 N-terminal tag to SENP1. Site directed mutagenesis by primer extension was used to make SENP1 mutants, C535S, C560S, C603S, C608S, C613S, C535S/C608S and C603S/C613S. In brief, two sets of overwrapping mutagenesis primers were used to amplify SENP1 catalytic domain using Phusion DNA polymerase. The SENP1 cDNA was cloned between Xba-I and BamH-I sites using Gibson assembly.

To produce wild type SENP1 without His×6 tag, human SENP1 catalytic domain cDNA was also cloned into immediately downstream of T7 promoter, between Nde-I and BamH-I of the pT7JLH plasmid, to avoid addition of a His×6 tag coding sequence.

For expression, Rosetta strain of *E. coli* (Novagen) was transformed with each expression vector and grown in 50 ml LB broth containing 100 µg/ml ampicillin at 37°C with shaking until A600 reached between 0.6 and 0.8. Each SENP1 protein was overexpressed by addition of 1 mM IPTG to the culture. After 2 hours induction, the *E. coli* pellet was harvested by centrifugation at 10,000 × g for 10 minutes. Inclusion bodies of SENP1 proteins were prepared from the pellet (*81*).

Each SENP1 protein with a His×6 tag was solubilized in 6 M guanidine HCl, 20 mM Tris, 0.1 % Tween20, 1 mM 2-mercaptoethanol, pH=8, and captured with a HisPur Nickel-NTA resin column (1.5 × 3cm). The column was washed with 4 M guanidine HCl, 20 mM Tris, 0.1 % Tween20, 1 mM 2-mercaptoethanol, pH=8 and then SENP1 protein was eluted with 4 M guanidine HCl, 50 mM sodium acetate, 0.05 % Tween20, pH=4. To remove Ni^2+^ leached out from nickel-NTA resin, the concentration of each SENP1 protein was adjusted to 4 mg/ml with 4 M guanidine HCl, 20 mM Tris, 0.1 % Tween20, 1 mM DTT, 50 mM EDTA and 0.2 M Tris, pH=8. Ten volumes of water were added to dilute the 4 M guanidine HCl to 0.4 M, which causes the precipitation of SENP1 protein. The precipitated SENP1 protein without nickel was recovered by centrifugation at 8000 × g for 20 minutes.

The inclusion bodies of wild type SENP1 without a His×6 tag, was further washed twice with 1 M guanidine HCl, 2% Triton X-100, 0.1M Tris, pH=8. The inclusion bodies were dissolved in 4 M guanidine HCl, 20 mM Tris, 0.1 % Tween20, 1 mM DTT, 50 mM EDTA and 0.2 M Tris, pH=8. The SENP1 without a His×6 tag was precipitated by addition of ten volumes of water. The SENP1 without a His×6 tag was recovered by centrifugation at 8000 × g for 20 minutes.

The precipitated SENP1 was dissolved in 6M guanidine HCl, 20mM Tris, 0.2M Arginine, 1mM DTT, pH=7.2 and the absorbance at 280 nm was measured to estimate protein concentration. SENP1 proteins were stored at −80°C until *in vitro* refolding. Refolding of SENP1 protein was carried out *in vitro* by gradual decrease of guanidine HCl concentration with dialysis (*82*) at 4°C. The concentration of SENP1 protein was adjusted to 1 mg/ml with 6 M guanidine HCl, 20 mM Tris, 0.1 % Tween20, 0.2 M arginine, pH=7.2, and for each refolding 4-5 mg of SENP1 was used. The cysteines of SENP1 proteins were reduced to sulfhydryl with 10 mM DTT for 1 hour at room temperature and dialyzed against 0.5 M guanidine HCl, 0.2 M arginine, 0.1 M Tris, 2 mM cysteine, mM cystine, 0.05 % Tween20, pH=7.2 for 24 hours, and then against the same solution without guanidine HCl for 24 hours. SENP1 proteins were refolded with/without 0.1 mM ZnCl_2_, and then ZnCl_2_ was removed by dialysis. Alternatively, SENP1 and mutants were prepared without ZnCl_2_ in the refolding buffer then incubated with 1 µM ZnCl_2_ for 1 hour to suppress SENP1 activities prior to reactivation by 0.1 mM GSH+10 µg/ml GRX1. SENP1 proteins were finally dialyzed against SENP1 assay buffer, 20 mM Tris, 0.1 M sodium chloride, 0.05 % Tween20, pH=7.2 for 24 hours. SENP1 proteins, used for the intracellular dialysis experiments, were prepared without 0.05% Tween 20. The final dialysis buffer was changed to fresh buffer twice. The precipitated SENP1 proteins were removed by centrifugation at 8000 × g for 10 minutes. Absorbance at 280 nm of the supernatant were measured to estimate the concentration of SENP1 proteins.

For the experiments to study the effect of divalent transient metals on SENP1 activity, 0.1 mM of CaCl_2_, CoCl_2_, NiCl_2_ and ZnCl_2_ was added in the refolding solution, and then removed by extensive dialysis against SENP1 assay buffer, 20 mM Tris, 0.1 M sodium chloride, 0.05% Tween20, pH=7.2

### Preparation of recombinant glutaredoxin 1, metallothinein 1X and thionein 1X

The coding sequence of human GRX1 cDNA(NM_002064.3/NP_002055.1, 1-106 residues) was cloned in the downstream of a His×6 tag coding sequence, between Xba-I and BamH-I sites of the pT7JLH plasmid (*83*). Full size human MT1X cDNA (NM_005952.4/ NP_005943.2, 1-61 residues) was cloned into immediate downstream of T7 promoter, between Nde-I and BamH-I sites of the pT7JLH plasmid (*83*), to avoid addition of a His×6 tag coding sequence. Recombinant GRX1 was produced in *E. coli*, purified and carried out *in vitro* refolding as described for recombinant SENP1 with a His×6 tag.

MT1X was expressed in Rosetta strain of *E. coli* (Novagen) as described above. The inclusion bodies of MT1X were washed twice with 2% Triton X-100, 0.1M Tris, pH=8, and then the inclusion bodies were dissolved in 6M guanidine HCl, 10 mM DTT, 10 mM EDTA and 0.2 M Tris, pH=8. The protein concentration was determined using BioRad Bradford protein assay (BioRad), and then adjusted to 1mg/ml. For preparation of MT1X or T1X, two milliliter of the solution (2mg MT1X) was used for each refolding with or without ZnCl_2_.

To prepare MT1X, which was loaded with zinc, the MT1X solution was exchanged to 6M guanidine HCl, 0.2M arginine, 0.5mM zinc chloride, 0.2M Tris, pH=7.2 using a centrifugal molecular cut off filter unit, Amicon Ultra-4 3kDa MWCO (Millipore), and then the final volume was adjusted to 4ml with the same solution. The MT1X solution was transferred into a Snakeskin 3.5kDa MWCO Dialysis tubing (Thermo Fisher), and then the dialysis was carried out at 4°C against three different solutions subsequently. The MT1X was dialyzed against 0.2M arginine, 0.1mM ZnCl_2_, 0.2M Tris, pH=7.2 for 24 hours, against 0.1mM ZnCl_2_, 0.1M Tris, pH=7.2 for 24 hours, and then 50mM Tris, pH=7.2 for 48 hours. The final dialysis buffer was changed to fresh buffer three times.

To prepare T1X, which is apoprotein of MT1X not carrying zinc, the volume of MT1X solution was adjusted to 4mL with 6M guanidine HCl, 10 mM DTT, 10 mM EDTA and 0.2 M Tris, pH=8, and then was transferred into a Snakeskin 3.5kDa MWCO Dialysis tubing. The refolding of T1X was carried out at 4°C against four different solutions subsequently; against 0.2M Arginine, 1mM EDTA, 1mM DTT, 0.2M Tris, pH=7.2 for 24 hours, against 1mM EDTA, 1mM DTT, 0.1M Tris, pH=7.2 for 24 hours, against 0.1mM DTT, 20mM Tris, pH=7.2 for 24 hours, and against 20mM Tris, pH=7.2 for 24 hours. The final dialysis buffer was changed to fresh buffer three times.

After refolding, both MT1X and T1X were concentrated using Amicon Ultra-4 3kDa MWCO to 200-300µg/mL, and the filtrates of Amicon Ultra-4 were kept for using as negative buffer control for MT1X and T1X assays, particularly to confirm the removal of free zinc in MT1X solution.

### In vitro SENP1-protease assays

Two assays were developed to measure SENP1 activity. Recombinant His×6-SUMO1-mCherry was used as a substrate for both assays. The cDNA of full size human SUMO1, including the cleavage site, and mCherry were amplified with PCR using Phusion DNA polymerase and the PCR fragments were cloned into a pT7JLH plasmid at Xba-I site using Gibson assembly to make pT7JLH-SUMO1-mCherry plasmid. Transformed Rosetta strain of *E. coli* was grown in 50 ml LB broth containing 100 µg/ml Ampicillin at 37°C with shaking until A600 reached between 0.6 and 0.8. His×6-SUMO1-mCherry protein was expressed by slow induction with 0.1 mM IPTG at 20°C overnight and the *E. coli* pellet was harvested by centrifugation at 10,000 × g for 10 minutes. Pink color of *E. coli* pellet indicated the proper folding of His×6-SUMO1-mCherry. The pellet was incubated with 500 µg/ml lysozyme, 1 mM PMSF, 5mM EDTA, 50 mM Tris, 0.2 M sodium chloride, pH=8 for 3 hours at 4°C, and then sonicated five times in short burst of 30 seconds on ice. The cell lysate was centrifuged at 10,000 × g for 10 minutes at 4°C, and the supernatant was dialyzed against 20mM Tris, 0.3M sodium chloride, pH=8. The soluble His×6-SUMO1-mCherry was purified with a HisPur Nickel-NTA resin column (1 × 5 cm). The purified His×6-SUMO1-mCherry was dialyzed against 5 mM EDTA, pH=8 for 6 hours and then SENP1 assay buffer (20 mM Tris, 0.1 M sodium chloride, 0.05 % Tween20, pH=7.2) for 24 hours. The final dialysis buffer was changed to fresh buffer twice. The concentration of His×6-SUMO1-mCherry protein was estimated by the absorbance at 280 nm.

For the first assay, HisPur Nickel-NTA resin was used to capture and precipitate the undigested substrate, and then the mCherry signal in the supernatant containing the digested substrate was measured. 100 nM of recombinant SENP1 protein was mixed with 1 µM of His×6-SUMO1-mCherry, and incubated at 37°C for 2 hours to cleave mCherry from His×6 tag-SUMO1. The undigested substrate was captured and removed by the addition of 25 µl of HisPur Nickel-NTA resin suspension (1:1, v/v). After vortexing for 10 seconds, HisPur Nickel-NTA resin was precipitated by centrifugation at 10,000 × g for 30 seconds at 4°C. Each supernatant was transferred into a 96 well clear bottom black plate and then fluorescent intensity, Ex544 nm/Em615 nm, was measured using Envision Multilabel plate reader (PerkinElmer) or Synergy HTX plate readers (Bio Tek Instruments) (**Fig. S2A**).

The second assay was used to examine the effect of MT1X and T1X on SENP1 activity. Because MT1X interfered the binding of Hisx6-SUMO1-mCherry to HisPur Nickel-NTA resin, native-PAGE (non-denaturing Tris/Glycine gel electrophoresis) was used to separate the digested substrate from undigested substrate. Recombinant SENP1 protein without His×6 tag (200nM) was mixed with 10 or 20 µg/mL (1.65 or 3.3 µM) MT1X or T1X, and 2.5µM His×6-SUMO1-mCherry in a 20µl reaction, incubated at room temperature for 45 minutes, and then separated by electrophoresis on non-denaturing Tris/Glycine gel. The mCherry signals of the undigested and digested substrate were detected using BioRad Gel Doc XR Imaging system (BioRad).

Because SENP1 activity changes during storage due to oxidation, the enzyme was fully activated prior to assays with 10 mM DTT at 4°C overnight. Amicon Ultra-4 10kDa MWCO was used to remove DTT and then replaced the buffer to SENP1 assay buffer, 20 mM Tris, 0.1 M sodium chloride, 0.05 % Tween20, pH=7.2. Final concentration of SENP1 and mutants were estimated by the absorbance at 280 nm. To preserve the activity of SENP1 25 % glycerol (w/w) was added and stored under nitrogen gas at −20°C.

### Analysis and statistics

Analysis was performed using FitMaster (HEKA Electronik) and GraphPad Prism (v7.0c). All data are shown as the mean ± SEM. Statistical outliers were identified and removed by an unbiased ROUT (robust regression followed by outlier identification) test. Normally distributed data were analyzed by the 2-tailed Student’s t test (for two groups), or ANOVA and Bonferroni post-test (for multiple groups). Data that failed normality tests were analyzed by the non-parametric Mann-Whitney test (for two groups), or the Kruskal– Wallis one-way analysis of variance followed by Mann-Whitney post-test (for multiple groups). A p-value less than 0.05 was considered significant. Zn^2+^Bind (https://zincbind.net/) was used to predict the Zn^2+^-binding site in SENP1 (2IYC) (*84*). Free Zn^2+^ for patch-clamp experiments was estimated with MaxChelator at https://somapp.ucdmc.ucdavis.edu/pharmacology/bers/maxchelator/.

## Acknowledgments

The University of Alberta is situated on Treaty 6 territory, traditional lands of First Nations and Métis people. We thank the Human Organ Procurement and Exchange (HOPE) program and Trillium Gift of Life Network (TGLN) for their work in procuring human donor pancreas for research, and James Lyon (Alberta) for his efforts in human islet isolation. We especially thank the organ donors and their families for their kind gift in support of diabetes research.

## Funding

National Institutes of Health NIH R01 GM132610 (WL)

Canadian Institutes of Health Research Foundation Grant 148451 (PEM)

Sino-Canadian Studentship from Shantou University (HL)

Canada Research Chair in Cell Therapies for Diabetes (ARP)

Canada Research Chair in Islet Biology (PEM).

## Author contributions

Haopeng Lin^1, 2, 8^†, Kunimasa Suzuki^1, 2^†, Nancy Smith^1, 2^, Xi Li^3^, Lisa Nalbach^4^, Sonia Fuentes^3^, Aliya F Spigelman^1, 2^, Xiaoqing Dai^1, 2^, Austin Bautista^1, 2^, Mourad Ferdaoussi^6^, Saloni Aggarwal^7^, Andrew R Pepper^7^, Leticia P Roma^5^, Emmanuel Ampofo^4^, Wen-hong Li^3^, Patrick E MacDonald^1, 2^ *

Conceptualization: HL, KS, ARP, EA, WL, PEM

Methodology: HL, KS, LN, LPR, ARP, EA, WL, PEM

Investigation: HL, KS, NS, XL, LN, SF, AFS, XD, AB, MF, SA, LPR

Visualization: HL, PEM

Supervision: ARP, LPR, EA, WL, PEM

Writing—original draft: HL, KS, PEM

Writing—review & editing: HL, KS, NS, XL, LN, SF, AFS, XD, AB, MF, SA, ARP, LPR, EA, WL, PEM

## Data and materials availability

All data are available in the main text or the supplementary materials.

