## Supplementary Figures and Table for "β-cell responses to high fat feeding: A role and mechanism for redox sensing by SENP1"

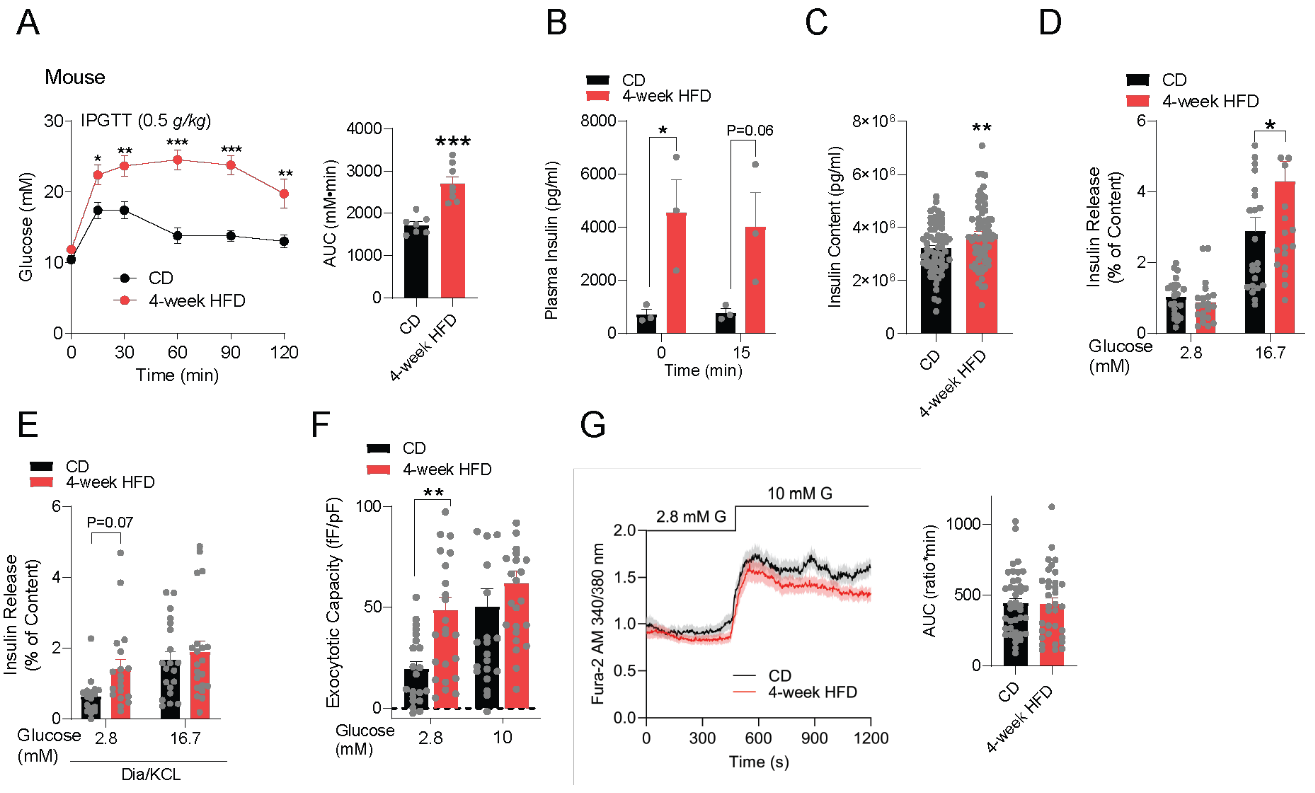


**Fig. S1. Increased insulin secretion and β-cell exocytosis after 4-week HFD**
**(A):** IPGTT of C57 mice after CD and 4-week HFD (n=7 and 8 mice). **(B):** Plasma insulin level during IPGTT (n=3 and 3 mice) **(C):** Insulin contents (n=22 and 22 mice) **(D):** Insulin secretion (n=8 and 8 mice) **(E):** Insulin secretion in the presence of 100 µM Diazoxide/ 30 mM KCL. (n=7 and 7 mice). **(F):** Exocytotic response (n= 21, 25, 23, 25 cells from 3 pairs of mice). **(G):** Single cell calcium response (n=40, 34 cells from 4 pairs of mice). Data are mean ± SEM and were compared with student t-test, one-way or two-way ANOVA followed by Bonferroni post-test. *$P < 0.05$, **$P < 0.01$.

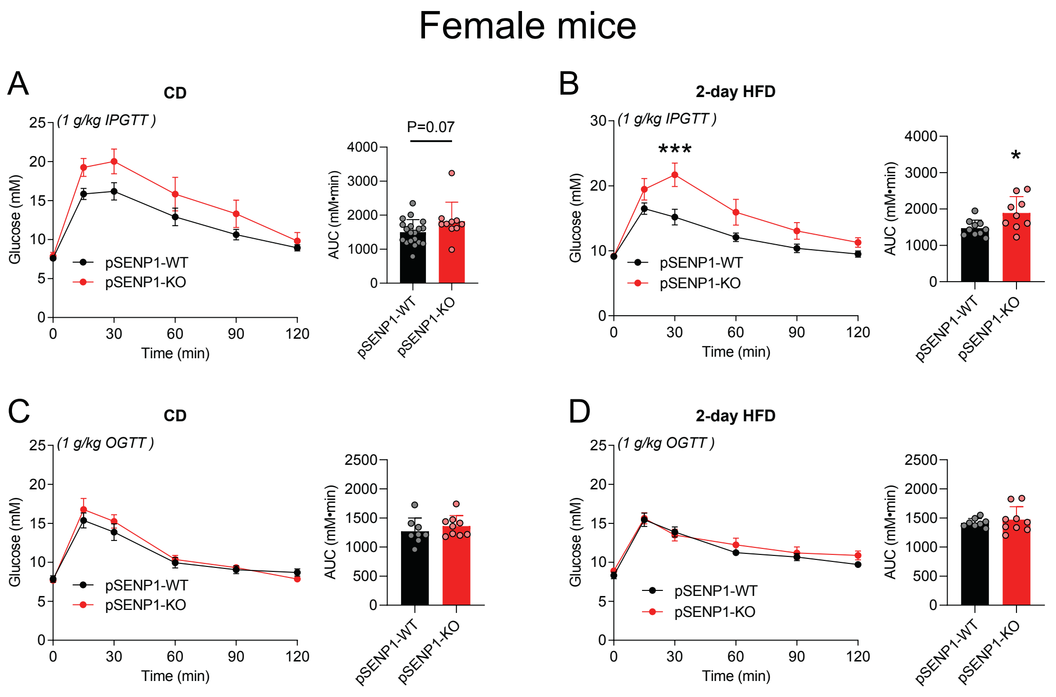


**Fig. S2. IPGTT and OGTT of female pSENP1-KO mice following HFD**
**(A-B):** IPGTT of female pSENP1-KO and pSENP1-WT mice fed CD or 2-day HFD (A - n=19, 10 mice, B - n=10, 9 mice). **(C-D)**: OGTT of female pSENP1-KO and pSENP1-WT mice fed CD or 2-day HFD (C - n=8, 9 mice, D - n=8, 9 mice). Data are mean ± SEM and were compared with student t-test or two-way ANOVA followed by Bonferroni post-test. **P* < 0.05, ***P* < 0.01.

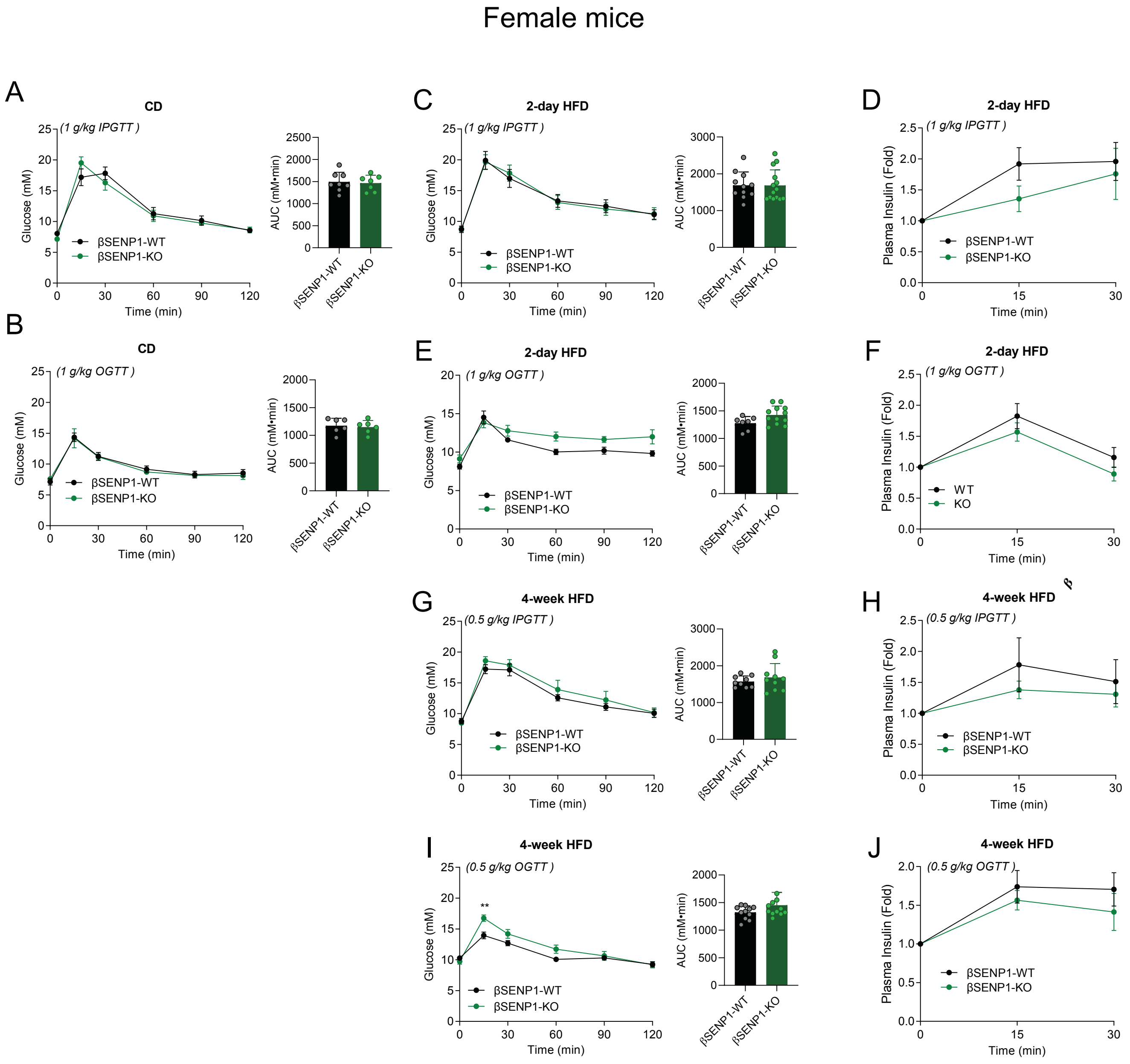


**Fig. S3. IPGTT and OGTT of female βSENP1-WT and βSENP1-KO following HFD**
**(A-C):** IPGTT following CD and 2-day HFD (A - n=8, 7 mice, B - n=11, 14 mice), and associated insulin secretion (C - n=8, 9 mice). **(D-F):** OGTT following CD and 2-day HFD (D - n=6, 6 mice, E - n=7, 12 mice), and associated insulin secretion (F - n=7, 8 mice). **(G-J)**: IPGTT and OGTT with plasma insulin level following 4-week HFD (G – n = 9, 10 mice, H - n=12, 11 mice, I - n=6, 6 mice, J - n=7, 8 mice). Data are mean ± SEM.

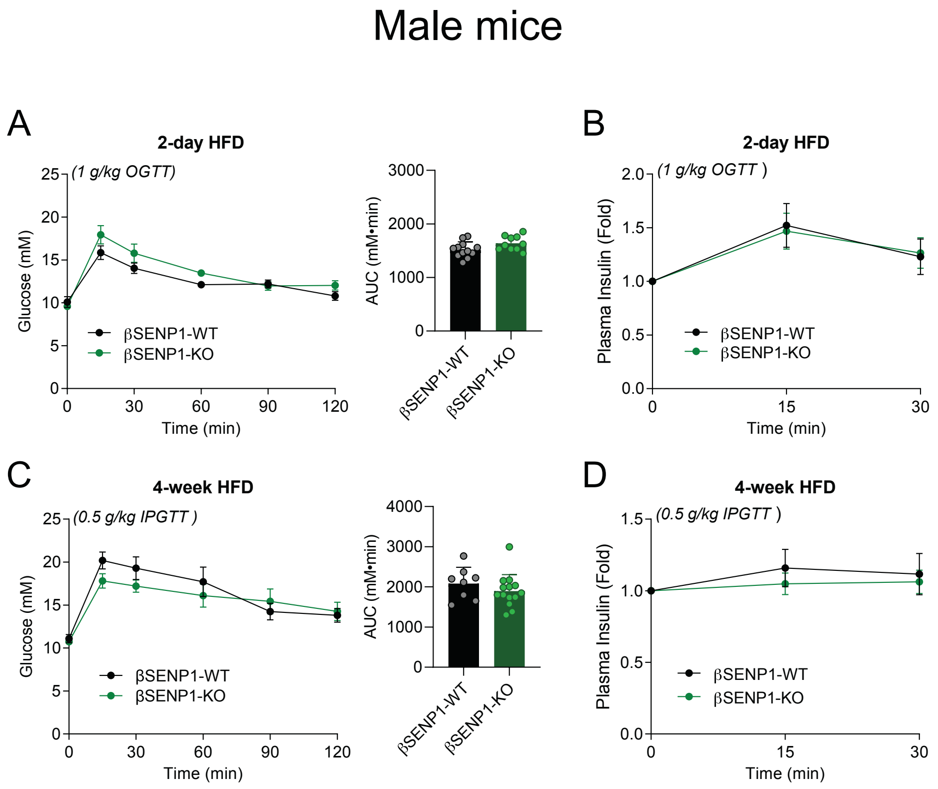


**Fig. S4. OGTT and IPGTT of male βSENP1-WT and βSENP1-KO following HFD**
**(A-B):** OGTT following 2-day HFD (A - n=12, 10), and associated insulin secretion (B - n=4, 6 mice). **(C-D):** IPGTT following 4-week HFD (C - n = 8, 14 mice), and associated insulin secretion (D - n=6, 10 mice). Data are mean ± SEM and were compared with student t-test or two-way ANOVA followed by Bonferroni post-test. **P* < 0.05, ***P* < 0.01.

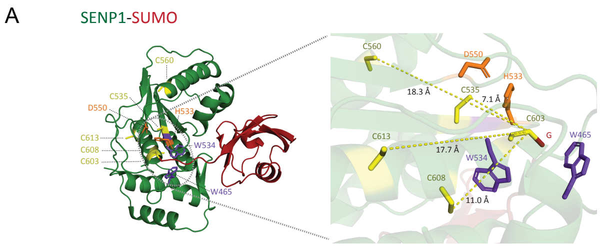


C

SENP1 WT

| Predicted Zinc-bind Site | Probability |
|---|---|
| Cys603, Cys535, His533 | 1 |
| Asp550, His533 | 1 |
| Asp468, His529 | 1 |
| Asp517, His491 | 1 |
| Asp529, His533, His539 | 1 |
| Glu430, His640 | 0.96 |

SENP1 C535S mutant

| Predicted Zinc-bind Site | Probability |
|---|---|
| Asp550, His533 | 1 |
| Asp468, His529 | 1 |
| Asp517, His491 | 1 |
| Glu430, His640 | 0.96 |

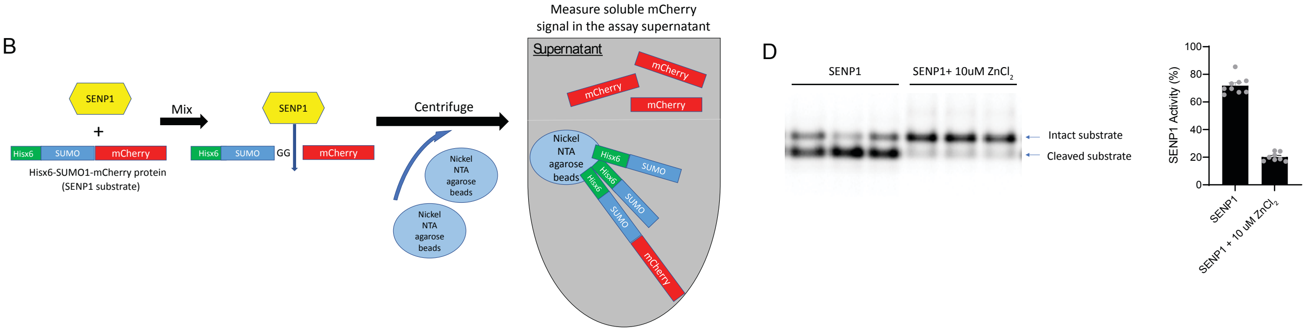


**Fig. S5. SENP1 activity in the presence of $Zn^{2+}$**

**(A):** Structure representation of SENP1-SUMO complex (ID: 2IYD, SENP1: green. SUMO: red) from PDB database. Potential redox-sensing cysteines (C535, C560, C603, C608, C613, yellow), another two catalytic triad (D550, H533, orange), and catalytic "floor" (W534, purple blue) and "lid" (W465, purple blue) were labeled. On the right panel, the catalytic region along with potential cysteines were outlined and zoomed in. Dashed lines indicated the measured distance (Å) between thiol group from catalytic (C603, yellow) to the ones from other potential cysteines (C535, C560, C603, C608, C613, yellow). **(B)**: Illustration of SENP1-activity assay with His×6-Nickle NTA purification. His×6-SUMO-mCherry protein is purified and incubated with SENP1 which cuts the specific sequence on SUMO protein, leaving the His×6-SUMO and mCherry fragment. Proteins with His×6 are removed with Nickel NTA agarose beads. The fluorescence of digested mCherry is measured and indicates the SENP1 activity. **(C):** Predicted $Zn^{2+}$ binding sites in SENP1. **(D):** Native-PAGE (non-denaturing Tris/Glycine gel electrophoresis) assay for SENP1-activity using substrate without His×6-Nickle NTA purification. Recombinant SENP1 protein without His×6 tag (200nM) was mixed with 10 µM $ZnCl_2$ and 2.5µM His×6-SUMO1-mCherry in a 20µl reaction, incubated at room temperature for 45 minutes, and then separated by electrophoresis on non-denaturing Tris/Glycine gel to visualized intact and cleaved substrate. SENP1 activity is calculated as the ratio of cleaved/intact substrate (n=9).

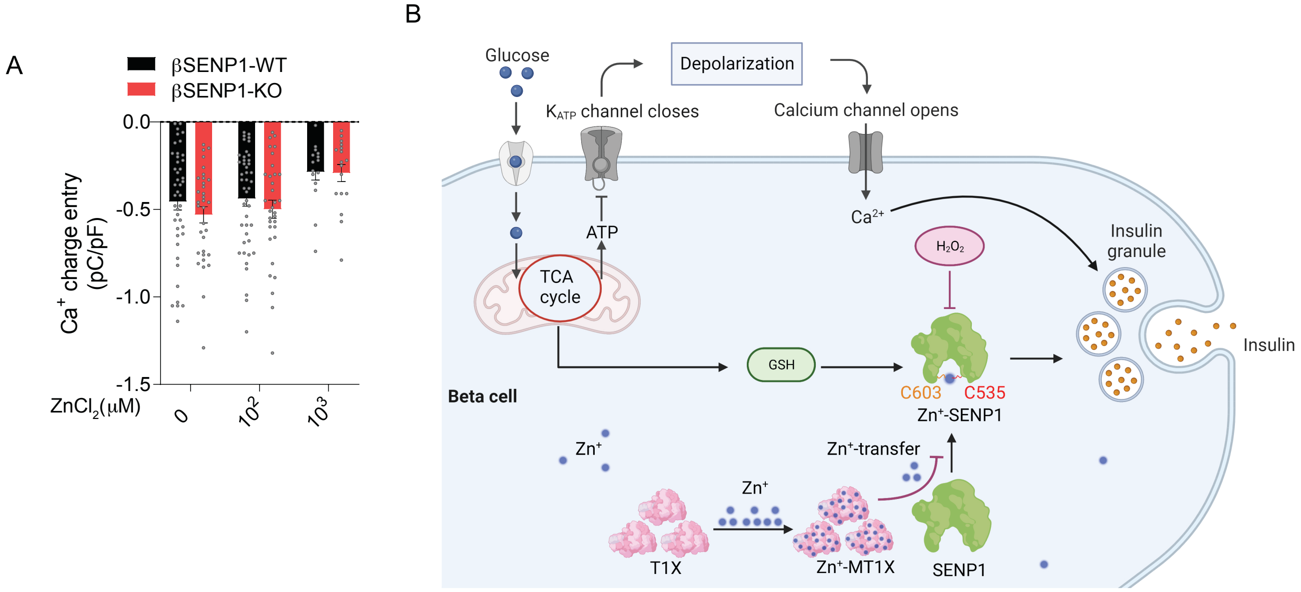


**Fig. S6. Effect of $Zn^{2+}$ on voltage dependent $Ca^{2+}$ entry in β-cells, and proposed scheme for SENP1 role in β-cell exocytosis, dependent upon redox and $Zn^{2+}$.**

**(A):** Effect of $Zn^{2+}$ on $Ca^{2+}$ charge entry during a 500 ms membrane depolarization from -70 to 0 mV in β-cells at 5 mM glucose from βSENP1-KO and βSENP1-WT mice (n= 23-46 cells from 6 pairs of mice). **(B):** Proposed scheme for SENP1 regulation by redox and $Zn^{2+}$ in β-cell exocytosis. $K_{ATP}$ – ATP-sensitive $K^+$ channel; TCA – tricarboxylic acid; ATP – adenosine triphosphate; GSH – reduced glutathione; SENP1 – sentrin specific SUMO protease; T1X – thionine; MT1X – metallothionine.

**Table S1. Human donors used in the study.**
Human islet donor information and detailed characteristics of each donor is available at www.isletcore.ca. ND: No Diabetes. T2D: Type 2 Diabetes.

| | | | | | | | | | | | | |
|---|---|---|---|---|---|---|---|---|---|---|---|---|
| ND | R012 | R039 | R050 | R051 | R055 | R058 | R060 | R061 | R062 | R067 | R072 | R073 |
| | R074 | R075 | R077 | R081 | R082 | R084 | R085 | R087 | R088 | R091 | R092 | R094 |
| | R096 | R097 | R098 | R099 | R100 | R101 | R102 | R104 | R105 | R106 | R108 | R109 |
| | R112 | R113 | R114 | R115 | R117 | R118 | R120 | R122 | R124 | R126 | R127 | R128 |
| | R130 | R134 | R135 | R136 | R138 | R139 | R140 | R142 | R144 | R145 | R146 | R147 |
| | R149 | R150 | R151 | R153 | R156 | R157 | R159 | R160 | R161 | R162 | R163 | R164 |
| | R165 | R168 | R169 | R172 | R174 | R175 | R176 | R179 | R180 | R181 | R182 | R183 |
| | R184 | R186 | R189 | R190 | R193 | R194 | R195 | R196 | R199 | R200 | R202 | R203 |
| | R204 | R205 | R208 | R210 | R215 | R216 | R217 | R218 | R220 | R221 | R223 | R226 |
| | R227 | R228 | R229 | R230 | R232 | R233 | R234 | R235 | R237 | R238 | R239 | R242 |
| | R243 | R245 | R246 | R248 | R250 | R251 | R252 | R253 | R254 | R255 | R256 | R260 |
| | R264 | R266 | R267 | R268 | R270 | R271 | R272 | R275 | R277 | R278 | R279 | R280 |
| | R282 | R284 | R285 | R286 | R288 | R290 | R291 | R292 | R294 | R299 | R301 | R302 |
| | R305 | R306 | R308 | R309 | R310 | R311 | R313 | R314 | R316 | R317 | R318 | R319 |
| | R321 | R322 | R323 | R324 | R326 | R327 | R328 | R330 | R332 | R333 | R334 | R335 |
| | R338 | R340 | R342 | R343 | R344 | R346 | R348 | R350 | R352 | R354 | R355 | R356 |
| | R358 | R360 | R361 | R363 | R364 | R365 | R366 | R367 | R368 | R369 | R370 | R372 |
| | R373 | R374 | R375 | R377 | R382 | R383 | R385 | R387 | R388 | R389 | R390 | R391 |
| | R392 | R393 | | | | | | | | | | |
| T2D | R031 | R064 | R070 | R071 | R083 | R107 | R110 | R125 | R131 | R143 | R152 | R154 |
| | R170 | R171 | R173 | R191 | R201 | R206 | R222 | R231 | R236 | R240 | R244 | R257 |
| | R263 | R265 | R307 | R312 | | | | | | | | |
